# Phylogeography of Ryukyu insular cicadas: Extensive vicariance by island isolation vs accidental dispersal by super typhoon

**DOI:** 10.1101/2020.12.10.419127

**Authors:** Soichi Osozawa, Kenichi Kanai, Haruo Fukuda, John Wakabayashi

## Abstract

Cicadas tend to be affected by vicariance reflecting poor mobility of nymphs underground and weak flying ability of adults. However, modern collection records of invasive cicada, combined with records of typhoon tracks, and newly obtained phylogeographic data suggest long distance, relatively instantaneous, dispersal of some vicariantly speciated cicadas. We address the importance of this typhoon dispersal mechanism applied to representative species of east Asian endemic cicadas of *Cryptotympana*, *Mogannia*, *Euterpnosia* and *Meimuna*. We combine BEAST-dated phylogenic and haplotype network analyses, modern collection data of non-native cicadas available in reports of the Japanese insect associations, modern typhoon records by Japan Meteorological Agency, and our own Quaternary geological constriction data. In conclusion, although Ryukyu endemic cicadas were vicariantly speciated, endemic cicadas on some islands were accidentally dispersed long distances to another island by typhoons, particularly those associated with super typhoons generated since 1.55 Ma.

## Introduction

In an earlier paper (Osozawa et al., 2017b), we showed extensive vicariant speciation acted on *Platypleura* cicada to generate each endemic population on each island of the Ryukyu chain, as well as the main islands of Japan and Taiwan. With a similar objective we analyzed the four cicada groups of *Cryptotympana*, *Mogannia*, *Euterpnosia* and *Meimuna*, which include endemic species and expected to find evidence of vicariance.

Contrary to expectation, *Platypleura kaempferi* is widespread in the Japanese islands, Korea and China, including the northern Tokara islands (see Fig. 1), and genetically similar (with the same molecular sequence; Osozawa et al., 2017b), suggesting a relatively strong dispersal ability of this cicada, including the ability to cross seaways.

**Fig. 1.**
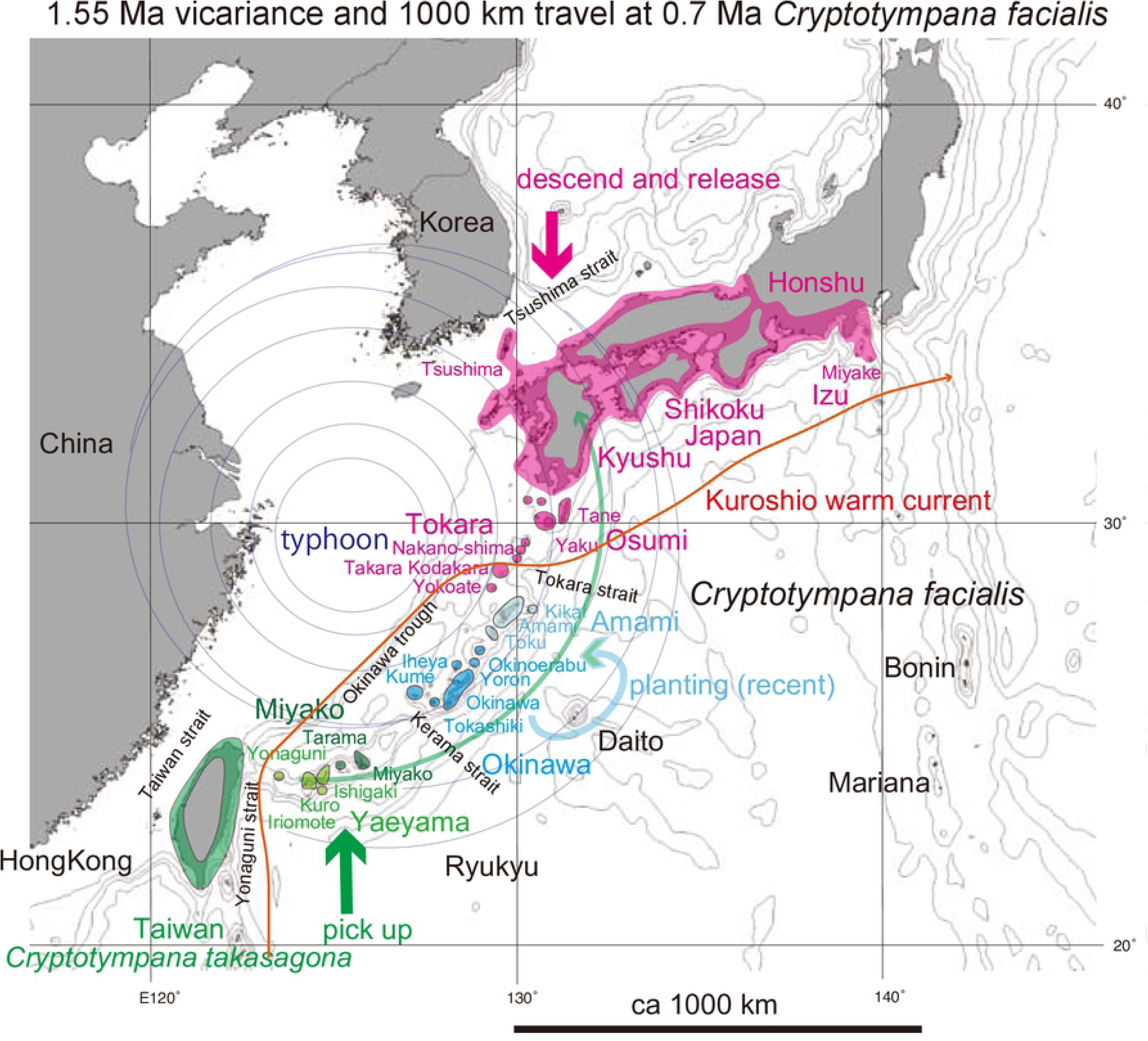
Distribution of *Cryptotympana facialis* in the Ryukyu and Japanese islands and *C. takasagona* in Taiwan. These species are lowland species as shown for the Japanese islands. Island populations are shown by color highlights. See Discussion; Green anticlockwise curved track: Super typhoon dispersal from Yaeyama (pale green) to Japan (red). The bold short arrows schematically (not to scale) show where the typhoon winds picked up cicada (green) and where the same cicada were released (red). Recent dispersal by human transport is also shown by the shorter counterclockwise path. The Kuroshio warm current is shown by the thin orange path with the arrow indicating northward to northeastward movement.

*Cryptotympana facialis* is known to have recently invaded the Amami islands, that originally lacked this cicada (Fukuda, 1991ab), and *Mogannia minuta* on the Okinawa main island recently extended its habitat (Sasaki, 2011). The former example interpreted to have resulted from artificial transplanting, probably from the Okinawa islands, whereas the cause of the latter factor was uncertain. From phylogeographic study, we may be able to detect the origin area of a cicada species using the extent of vicariant speciation. Recognition of the cicada source may give insight into the mechanism of dispersal, being it an artificial or natural mechanism.

Cicada is commonly absent from oceanic islands, but exceptions include *Euterpnosia chibensis daitoensis* on the Daito islands, *Meimuna boniensis* on the Bonin islands, and *Meimuna opalifera* on the Izu islands (island localities in Fig. 1). A phylogeographic study for such cicadas may help ascertain the origin, date of dispersal, and dispersal mechanism.

We have shown that range fragmentation acted severely on these island cicada groups, but super typhoons generated since ca.1.55 Ma (Osozawa & Wakabayashi, 2015) played an important role in long distance dispersal to oceanic islands. We also found evidence of intra continental island dispersal such as for *Cryptotympana facialis*. Dispersal of species by typhoon has not been adequately addressed as a dispersal mechanism to date. In this paper we consider multiple data threads, including recent typhoon track records, meteorology and physics, cicada ecology and collection records, transplantation records, and also formative (geologic) history of islands terraces.

## Materials

### Taxon sampling and the analytical aims

We collected four cicada groups from the Ryukyu islands, as well as from the Japan islands, Taiwan, and Chinese mainland (Table 1). See Japanese specimen photos in Hayashi & Saisho (2011), and Taiwan specimen photos in Chen (2011).

**Table 1.** Cicada species collected and analyzed in this paper.

#### Cryptotympana (Fig. 1)

We collected *Cryptotympana takasagona* and *C. holsti* from Taiwan, *C. mandarina* from Hong Kong, and *C. atrata* from China and Korea. Sequence data for *C. consanguinea* in Philippine and *C. mandarinais* in Vietnam are available in GenBank/DDBJ.

*C. facialis* is endemic in the Japanese main islands as well as the Ryukyu islands, northern Izu islands, and Tsushima island (Fig. 1). Its song and habitat are similar to *C. takasagona* in Taiwan according to Hayashi & Saisho (2011). A white band along the margin of abdominal segment III is a characteristic of the Yaeyama islands population and Tarama island specimens. This band especially broad on Yonaguni island specimens, but completely absent from Okinawa and Miyako island specimens. This white band is narrow or absent for species inhabiting the Japan main islands, but somewhat broad for the Osumi and Tokara island populations. Emergence time tends to early for the Yaeyama islands population situated to the south, and delayed for the more northerly Japanese population, but exceptionally delayed for the Iriomote island population (a part of the Yaeyama islands population). These morphological and ecological characters may be related to the vicariance on these islands, suggesting the potential utility of phylogenetic study.

*C. facialis* inhabits lowland regions including coastal area less than ca. 100 m altitude (red area in Fig. 1 for the Japan main islands) such as the Ryukyu islet areas. *C. facialis* was originally absent from the Amami islands of Amami Oshima, Toku, and Kikai (Fukuda, 1991ab; Fig. 1), possibly as a consequence of the geologic evolution of these islands (see Discussion). *C. facialis*, however, has been found on these islands since 1991, and has been proposed to have arrived as a result of artificial dispersal from Okinawa beginning in 1989. The newly found Amami islands *C. facialis* lack the white abdominal band similar to the Okinawa island population. We will show that *C. facialis* collected from the Amami islands is genetically common to those in the Okinawa islands.

#### Mogannia (Fig. 2)

*Mogannia minuta* is distributed on Okinawa island, and the Miyako and Yaeyama islands. Although this species was first reported from southern Taiwan and described (Matsumura, 1907) and a specimen (11-15 June, 1937, Heng-chun, M. Chujo leg.) was examined by Hayashi (1976), no record was known from Taiwan after World War II, so we could not include the Taiwan specimen in our analyses.

The emergence time of *M. minuta* is late March to late June with a peak of late April for the Yaeyama islands, end of April to May for the Miyako islands, and May for the Okinawa islands, although some cicadas continue to sing until June in Yagachi-jima (new habitat; Fig. 2 inset) of the northern Okinawa islands. This variation of emergence may be related to the vicariance in these islands.

**Fig. 2.**
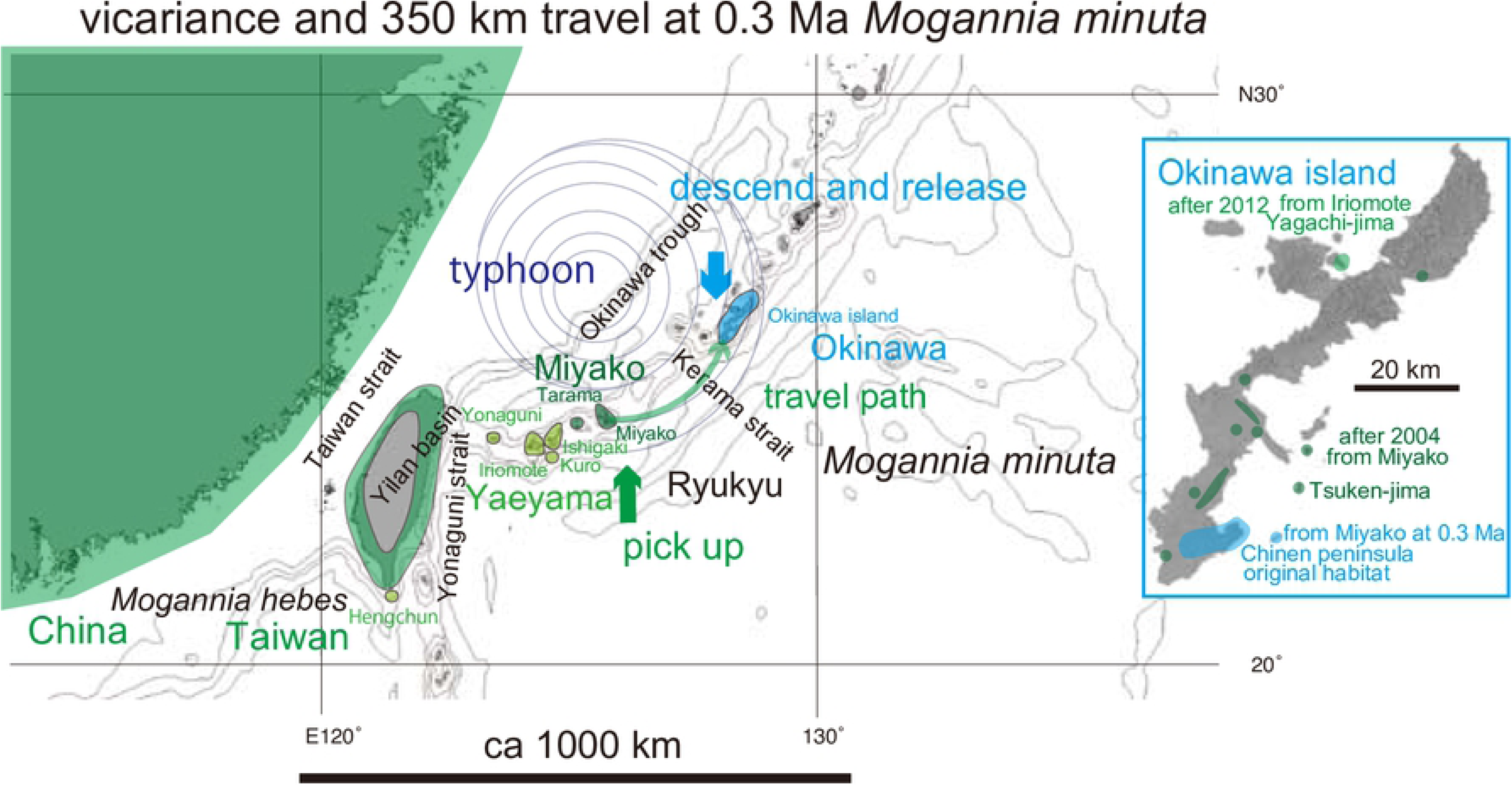
Distribution of *Mogannia*. Island populations are shown by color highlights. See Discussion; Green anticlockwise curved track: Typhoon dispersal at 0.3 Ma from Miyako (green) to Okinawa (blue). The bold short arrows schematically (not to scale) show where the typhoon winds picked up cicada (green) and where the same cicada were released (blue). Inset: Original habitat in the southern Okinawa and new habitats (blue). We clarified that only the Yagachi-jima population (green) has the same COI sequence with the southern Iriomote-jima, Yaeyama, and dispersed from there.

Habitats of *M. minuta* on Okinawa island were originally restricted to the southern part of the Chinen peninsula (Fig. 2 inset). However, Sasaki (2011) reported 2004 reports of this species from north of the Chinen peninsula, up to 20 km from the former area of distribution. Internet records dating from 2012 were found for documenting the presence of cicada in northern Okinawa, including Yagachi-jima (Fig. 2 inset). Our study is designed to test the mechanism of habitat expansion of this species. Note that nymphs of this species are usually underground for two years (Hayashi, 1976), so dispersed female(s) at a new location should have egged two years before the first records of emergence and singing.

*Mogannia hebes* is known from Taiwan and southern China (Fig. 1), and we collected specimens for comparison with *M. minuta*. Note that the Yilan basin and Lanyang valley (the on-land continuation of the Okinawa trough on Taiwan) constitute a physical biological barrier separating the northwestern (Chinese side) and southeastern (Ryukyu side) of Taiwan (Fig. 2; Osozawa et al., 2017c).

#### Euterpnosia (Fig. 3)

*Euterpnosia* species are distributed from the Himalaya to southern China, Taiwan, Ryukyu islands, and Japan. It diversified into 13 species in Taiwan. In the Ryukyu islands, three endemic species are known from the Amami, Okinawa, and Yaeyama islands, whereas *E. chibensis daitoensis* is known from the Daito oceanic islands (Fig. 3) off shore (east) of the Ryukyu islands. We will address this apparently anomalous distribution with the analyses in this paper.

**Fig. 3.**
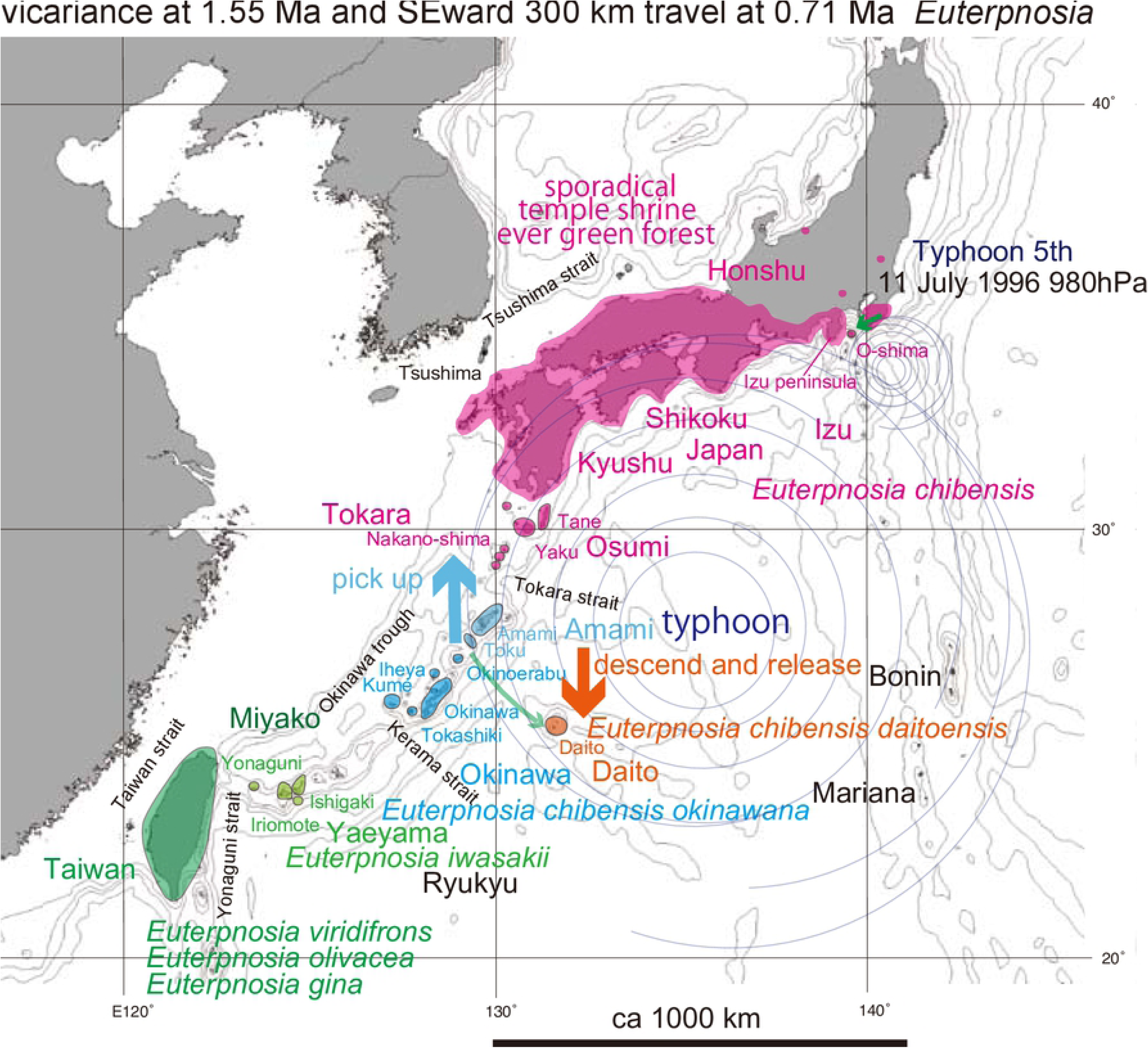
Distribution of *Euterpnosia*. Island populations are depicted by color highlights. See Discussion; Green anticlockwise curved tracks: Typhoon dispersal from Toku to Daito islands (SW part of map), and also from Honshu to O-shima (much shorter path on NE part of map). The bold short arrows schematically (not to scale) show where the typhoon winds picked up cicada (red) and where the same cicada were released (orange).

In Japanese main islands, *E. chibensis chibensis* is restricted to evergreen forests of southwest Japan, particularly protected forests associated with Japanese shrines. Three isolated habitats constitute the northern limit of this species within Honshu (three small red circles in Fig. 3), and the isolated distribution pattern suggests progressive vicariant speciation. In contrast, *E. chibensis chibensi*s was also recorded from the Oshima islet, northern end of the Izu oceanic islands, 20 km from the Izu peninsula (Fig. 3), since 1998. This may reflect and a similar but a smaller-scaled dispersal process comparable to that which may have populated the Daito islands.

#### Meimuna (Figs. 4 and 5)

*Meimuna kuroiwae* has a limited distribution on the Okinawa, Amami, Tokara, and Osumi islands, and at Cape Sata of southernmost Kyushu, and is absent on the Yaeyama islands. Local morphological variation is recognized (Hayashi & Saisho, 2011), and vicariant speciation is expected.

*Meimuna boniensis* is known from the Bonin islands, with a distance of 1000 km from the Okinawa islands and also from Tokyo (Fig. 4). The morphology and song is similar to *M. kuroiwae*, and considered to have recently derived from some of the above localities by transplanting (Hayashi & Saisho, 2011). Because *M. boniensis* is rigidly and nationally protected, it is usually impossible to analyze, but the phylogenetic tree was informally shown (Nagata, 2019). Although he did not upload the sequence data into GenBank/DDBJ, *M. boniensis* was represented as a sister of *M. kuroiwae* in Okinawa, and unrelated to the historic transplanting. Therefore, *M. boniensis* may have naturally dispersed from Okinawa to Bonin in pre historic time.

**Fig. 4.**
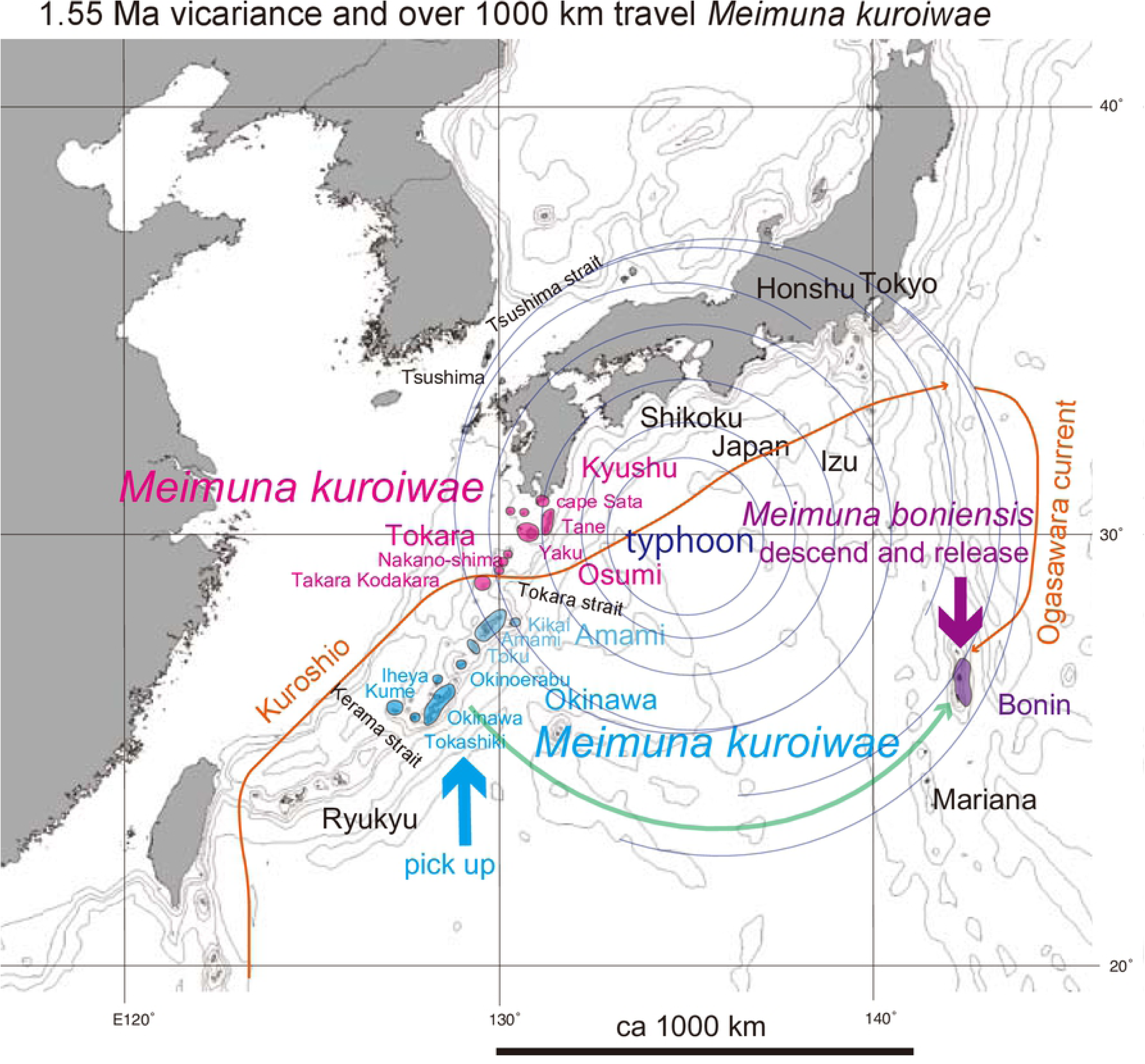
Distribution of *Meimuna kuroiwae*. Island populations are shown by colored highlights. See Discussion; Green anticlockwise curved track: Super typhoon dispersal from Okinawa (blue) to Bonin (purple, *Meimuna boniensis*). The bold short arrows schematically (not to scale) show where the typhoon winds picked up cicada (blue) and where the same cicada were released (purple). The Kuroshio warm current is shown by the thin orange path, and the southward Ogasawara current is also shown.

We also analyzed *Meimuna opalifera*, cosmopolitan in Japan, Taiwan, Korea, and China (Fig. 5), although this species is lacking and replaced by *M. oshimensis* and *M. iwasakii*, endemic in the Ryukyu islands (the latter is also known in Taiwan). The cicada’s song on the Osumi islands, Taiwan, and Korea is distinct from that in the Japanese main islands (Hayashi & Saisho, 2011), and this may reflect genetic differences so we analyzed this species. This species is also known from the Hachijo-jima and Aoga-shima (southern limit) islands of the Izu oceanic islands, 300 km apart from Tokyo (Fig. 5). We noted that the cicada’s song on Hachijo-jima was different from that of the Japanese main islands, so we added a Hachijo specimen to our analyses.

**Fig. 5.**
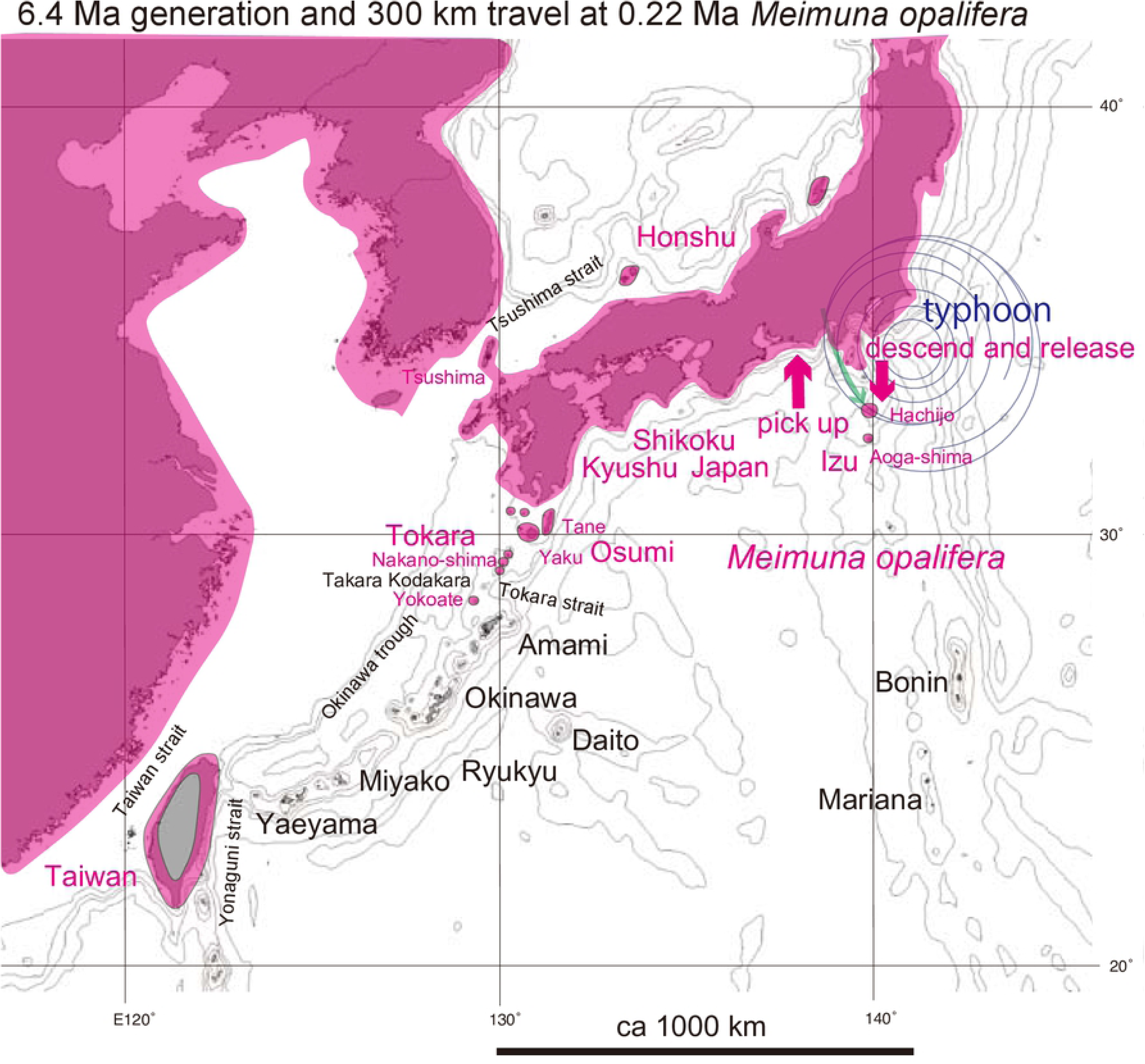
Distribution of *Meimuna opalifera*, that is absent in the Ryukyu islands. See Discussion; Green anticlockwise curved track: Typhoon dispersal from Honshu to the northern Izu islands of Hachijo and Aoga-shima. The bold short arrows schematically (not to scale) show where the typhoon winds picked up cicada (red) and where the same cicada were released (red).

### Analytical methods

#### Applied DNA Sequence

Mitochondrial COI and nuclear 18S rRNA sequence data from our 92 collected specimens are presented in Table 1. Primers used, amplifications, and sequencing are given in Osozawa et al. (2017). These sequences were aligned by ClustalW in MEGA X (Kumar et al. 2016). The COI sequence data comprise 1,534 bp, and the 18S rRNA sequence 874 bp, and the resolution to construct phylogenetic tree and haplotype network was sufficient, as we experienced in the *Platypleura* paper (Osozawa et al., 2017).

#### Phylogenetic analyses associated with fossil and geological event calibrations by BEAST v1.X

A Bayesian inference (BI) tree (Fig. 6) was constructed using the software BEAST v.1X (Suchard et al., 2018), running BEAUti, BEAST, TreeAnnotator, and FigTree, in ascending order. Before operating the BEAST software, the BEAGLE Library must be downloaded.

**Fig. 6.**
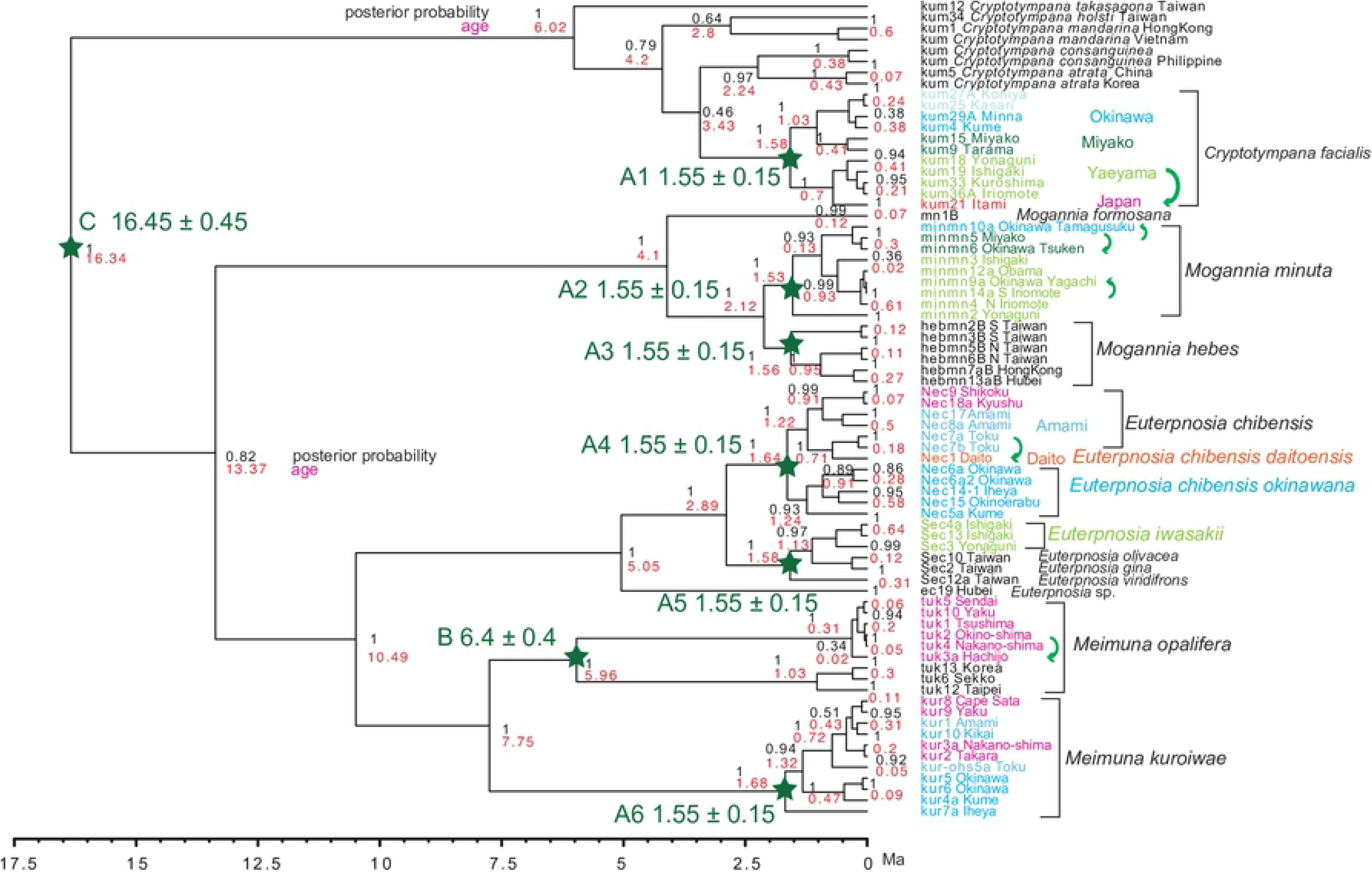
BI tree for *Cryptotympana*, *Euterpnosia*, *Mogannia*, and *Meimuna*. Vicariantly speciated island populations are shown by different colored text labels. Calibration points and dates are shown by green stars and corresponding large font green text labels. Black numbers on each node: posterior probability, red numbers: age (in Ma). The green curved arrows depict dispersal after the vicariance.

For graphic explanation of the operation of this software, see the “BEAST operating manual” at: http://kawaosombgi.livedoor.blog/archives/11386037.html

Calibrations points are shown on Fig. 6, and these dates were input in “Priors” in BEAUti; they are summarized below. Corresponding ingroup species were included in ingroup taxa by “Taxon Set” on the “Taxa” screen in BEAUti.

Calibration point A (A1 to A6) is after our geological event calibration that adopts a 1.55 Ma date based on multiple biostratigraphic and supporting radio-isotopic dates connected to various geologic relationships in the Ryukyu islands region (Osozawa et al., 2012). This geologic event calibration was also used in a previous study of *Platypleura* cicadas by our group (Osozawa et al. 2017).

Calibration point B: Crown *Meimuna opalifera*: Fossil *M. protopalifera* was found from the Itamuro Formation, Tochigi, Japan (Fujiyama 1982; Yoshikawa 2005), and the fission track age of the correlative terrestrial strata of the Nashino Formation, Sendai, is 6.4 ± 0.4 Ma (Fujiwara et al. 2008).

Calibration point C: Crown *Cryptotympana*: Fossil *C. incasa* and *C. miocenica* were found from Shanwang, Shandong, China (Moulds, 2018), and these strata are considered to be time correlative to the European MN5 mammalian stage (16.45 ± 0.45 Ma; Roček et al. 2011).

#### Haplotype network analyses by Network

In DnaSP 6.12.13 (Rozas et al., 2017), sequence files such as fasta files were converted into rdf files.

In the platform software of Network 10, inputting the above rdf file, calculation and graphic outputting was performed.

For graphic explanation of the operation of these softwares, see the “Network tutorial” at: http://kawaosombgi.livedoor.blog/archives/21400217.html

## Results

### BI tree (Fig. 6)

The BI tree consists of the *Cryptotympana*, *Mogannia*, *Euterpnosia,* and *Meimuna* clades. The *Cryptotympana* clade consists of *C. facialis* and the remaining five species, and *C. facialis* is not a sister of *C. takasagona*. The *Meimuna* clade consists of *M. opalifera* and *M. kuroiwae* clades. *Cryptotympana facialis*, *Mogannia minuta*, *Mogannia hebes*, *Euterpnosia chibensis*, *Euterpnosia iwasakii* + Taiwan *Euterpnosia*, *Meimuna kuroiwae* were simultaneously differentiated at 1.55 Ma as calibrated by this date.

For *Cryptotympana facialis*, note that the sequence data of kum25 Kasari (Amami) = kum26 Naze (Amami) = kum10 Toku Amagi = kum28A Toku Amagi = kum28B Toku Amagi = kum2 Yoron = kum30 Okinawa Kunigami = kum31 Okinawa Nago = kum20 Okinawa Yaese (Table 1), and the sequence data are common on the Okinawa main island and the Amami islands, except for kum27AB Koniya. The Okinawa (+ Amami) population is a sister of the Miyako population, but the northernmost Japan (+ Osumi and Tokara) population is a sister of the southernmost Yaeyama population (Fig. 1).

For *C. facialis*, note that kum21 Itami = kum14 Kagoshima = kum23A Tane = kum22A Yaku = kum8 Nakano-shima = kum13 Takara = kum24A Kodakara (Table 1), and the sequence data are regionally the same in the Japanese main islands and the Osumi and Tokara islands. The COI-5P sequence data found in GenBank/DDBJ are the same as our data for a total of 106 specimens collected from Honshu and Kyushu, although four specimens differ one base pair relative to these 106 specimens.

For *Mogannia minuta*, sequence data of minmn9a Okinawa Yagachi (Fig. 2 inset) is identical to minmn14a S Iriomote (Yaeyama). The sequence data of minmn6 Okinawa Tsuken (new habitat of Okinawa-jima; Fig. 2 inset) is similar to minmn5 Miyako, but more differentiated for minmn10a Okinawa Tamagusuku (specimen collected at the original habitat in southern Okinawa-jima; Fig. 2 inset).

For *Mogannia hebes*, the northern Taiwan population is a sister of the Chinese mainland population west of the Taiwan strait, and the southern Taiwan population is a sister of northern Taiwan + China population.

For *Euterpnosia chibensis*, N-ec9 Shikoku differs by only one base pair from N-ec18a Kyushu, and this Japan population is a sister of the Amami population. Nec1 Daito (*E. chibensis daitoensis*) is a sister of Nec7ab Toku, and these have a sister relationship to the Japan + Amami population. *E. chibensis* in these areas is a sister of *E. chibensis okinawana* on Okinawa. *Euterpnosia iwasakii* (Yaeyama) is morphologically similar to *E. viridifrons* (Taiwan), but a sister of *E. olivacea* + *E. gina*.

For *Meimuna kuroiwae*, the northern population including Cape Sata, Osumi, Tokara, and Amami islands, is a sister of the southern population of Okinawa, but kur7a Iheya is strongly differentiated.

*Meimuna opalifera* on the Japan islands and northern Tokara islands was mildly or not differentiated within these areas, but moderately differentiated on the Hachijo island relative to the others (= sister of the Japan population). The Japan population including the Hachijo island is a sister of the Korea + China + Taiwan populations, and tuk12 Taipei is strongly differentiated relative to the others. *M. opalifera* is absent from the main part of the Ryukyu islands.

### Haplotype network

#### Cryptotympana (Fig. 7)

The six analyzed species contain many genetically distinct interspecies haplotypes, and *Cryptotympana facialis* is genetically distinct from *C. takasagona*. Species level differentiation is recognized in *C. atrata* in Korea and China, *C. mandarina* in Hong Kong and Vietnam, and *C. consanguinea* on Mindanao of the Philippines.

Three areal haplotypes of the Okinawa, Miyako, and Yaeyama islands, from north to south, respectively, are recognized for *Cryptotympana facialis*, and these haplotypes, separated by major straits such as the Kerama strait (Fig. 1). The northernmost Japanese haplotype, however, is included in the southernmost Yaeyama haplotype, as is not an independent haplotype. In addition, the Japanese haplotype kum21 Itami contains the same base pair haplotype as the Osumi and Tokara haplotypes and kum22A Yaku and kum24A Kodakara + 102 haplotypes of Honshu and Kyushu, and the 4 haolotypes that differ by one base pair (red colored region in Fig. 1).

#### Mogannia (Fig. 8)

A haplotype network comprises *Mogannia minuta* and *Mogannia hebes*. *Mogannia formosana* is a distinct haplotype.

The *Mogannia minuta* network consists of Yaeyama and Miyako-Okinawa haplotypes. An expected Taiwan haplotype is shown on this Fig.. A haplotype consists of the same sequence as min-mn14a S Iriomote and min-mn9a Okinawa Yagachi (Fig. 2 inset). In the Miyako-Okinawa network, min-mn5 Miyako is similar to min-mn6 Okinawa Tsuken, and min-mn10a Okinawa Tamagusuku (original habitat).

The *Mogannia hebes* network consists of Chinese, northern Taiwan, and southern Taiwan haplotypes, differentiated from one another.

#### Euterpnosia (Fig. 9)

Haplotype network consists of northern Japan-Amami-Okinawa and southern Yaeyama-Taiwan haplotypes. The Chinese haplotype is distinct from the remaining haplotypes.

*Euterpnosia chibensis daitoensis* of ec1 Daito is close to *E. chibensis* of N-ec7ab Toku. *E. chibensis* haplotypes as a whole comprise a network, whereas the Japanese haplotypes of N-ec9 Shikoku and N-ec18a Kyushu constitutes a distinct network from the Amami network. E*. chibensis okinawana* also constitutes a distinct network.

S-ec3 Yonaguni of *E. iwasakii* is close to S-ec12a of *E. viridifrons*, but S-ec2 of *E. gina* and S-ec10 of *E. olivacea* constitute a single Taiwan network with S-ec12a.

#### Meimuna (Fig. 10)

*Meimuna kuroiwae* network consists of the Cape Sata-Osumi-Tokara-Amami and Okinawa haplotypes. Haplotype of southern kur10 Kikai is connected with kur9 Yaku with one base substitution, and then northernmost kur8 Cape Sata with four base substitutions. The expected *Meimuna boniensis* haplotype is connected with the Okinawa haplotype, according to Nagata (2019).

*Meimuna opalifera* network consists of the Japan and Korea-China-Taiwan haplotypes. The Japanese haplotypes are not differentiated, and a haplotype consists of the same sequence of tuk1 Tsushima, tuk2 Okino-shimaa, and tuk4 Nanano-shima. However, tuk3a Hachijo is differentiated relative to the remaining Japan haplotypes.

## Discussion

### Vicariance due to separation by seaways

In Fig. 6, *Cryptotympana facialis*, *Mogannia minuta*, *Mogannia hebes*, *Euterpnosia chibensis*, *Euterpnosia iwasakii* + Taiwan *Euterpnosia*, *Meimuna kuroiwae* were simultaneously differentiated at 1.55 Ma, which suggests vicariant speciation acted on the most recent common ancestors of these ingroup species. As we showed for *Platypleura* cicada (Osozawa et al., 2017b), the differentiation trigger was the physical separation of islands from the Chinese mainland as a result of with rifting (by sea-floor spreading) of the Okinawa trough that started at 1.55 Ma (Osozawa et al., 2012).

In the haplotype network of *Cryptotympana facialis* (Fig. 7), *Mogannia minuta* and *Mogannia hebes* (Fig. 8), *Euterpnosia chibensis* and *Euterpnosia iwasakii* + Taiwan *Euterpnosia* (Fig. 9), and *Meimuna kuroiwae* (Fig. 10), red double line indicates vicariance between island groups within a single species (whereas *E. iwasakii* + Taiwan *Euterpnosia* is an extra species level), although orange line indicates species level classification with the above exception. Such networks separated by red double lines in these Fig.s are geographically separated by major seaways also shown in Fig.s 1 ∼ 4. Classification into each haplotype in these haplotype networks (Fig.s 7 ∼ 10) reflects milder vicariance separated by minor seaways, although sporadically connected with each other during periods of lower sea level during glacial episodes (cf., Osozawa et al., 2017b).

**Fig. 7.**
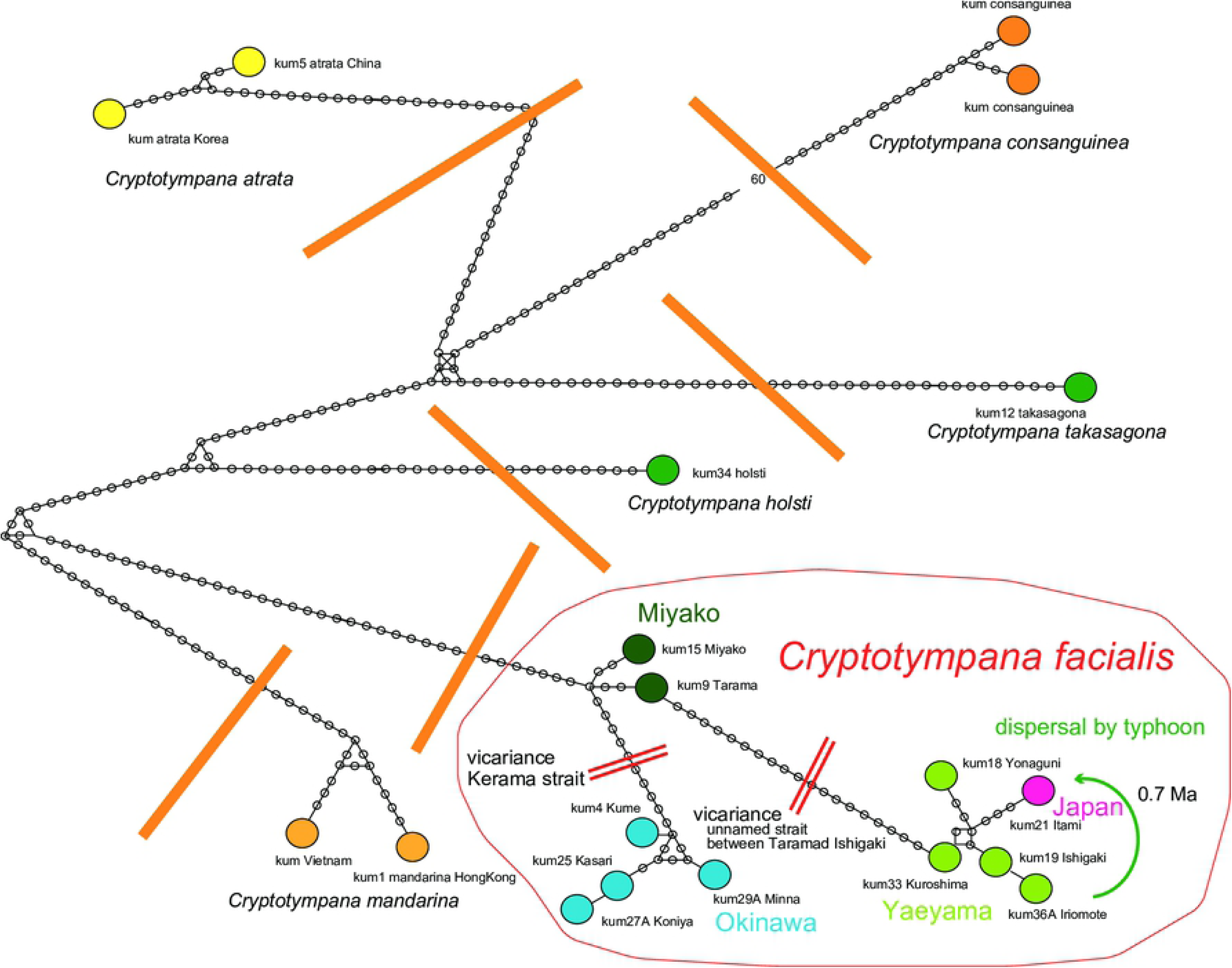
Haplotype network of *Cryptotympana. C. facialis* was vicariantly speciated due to isolation of island groups of Okinawa, Miyako, and Yaeyama, separated by major straits (red double line) formed since 1.55 Ma, and three island group haplotypes or populations were formed. However, the Japan haplotype was included in the Yaeyama haplotype network, suggesting a long-distance dispersal (green curved arrow) from Yaeyama (southern end of Ryukyu) to Japan at 0.7 Ma (Fig. 6), crossing Miyako and Okinawa. Orange heavy line: *C. facialis* is separated by the other *Cryptotympana* species including *C. takasagona*.

**Fig. 8.**
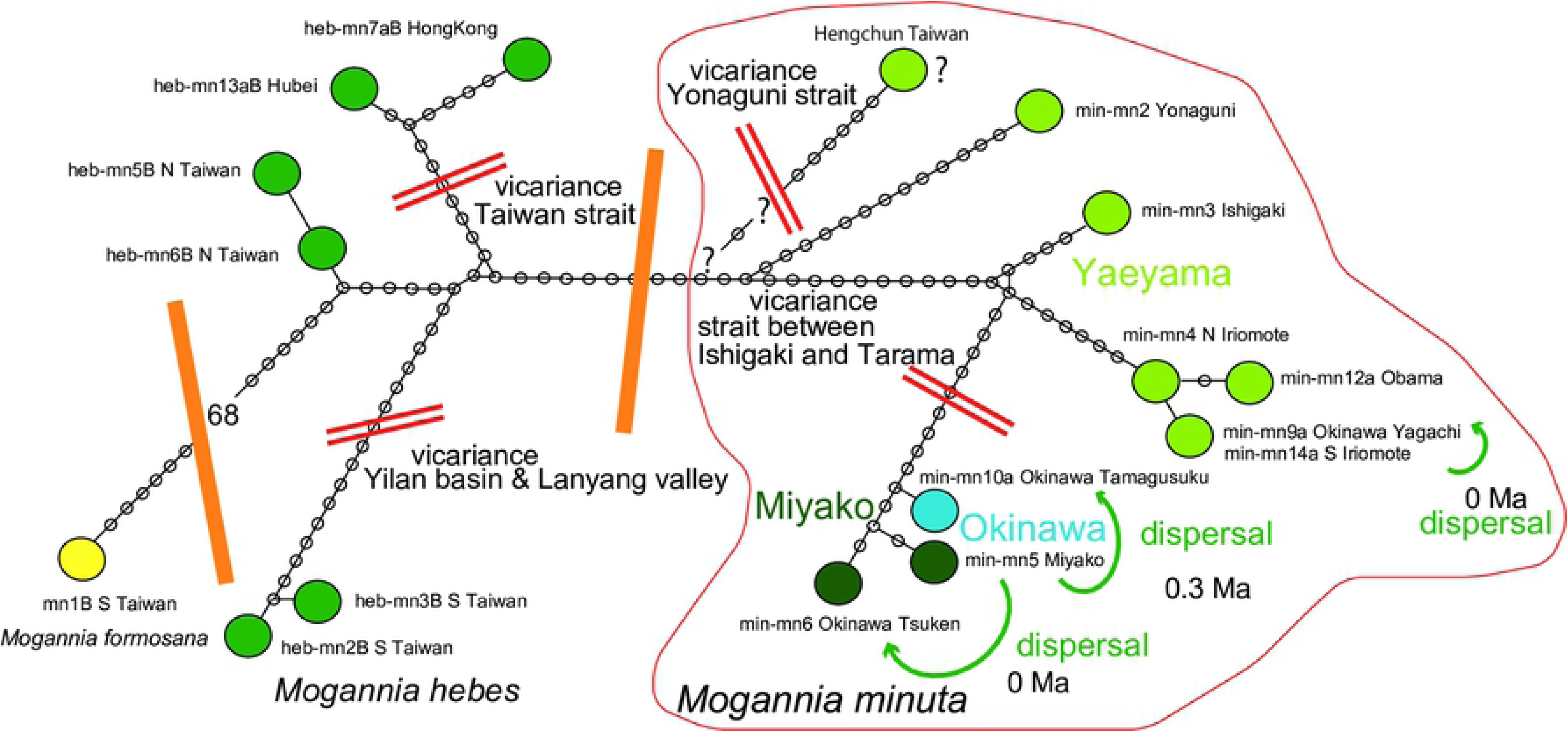
Haplotype network of *Mogannia.* On each of Yaeyama islands, *Mogannia minuta* constitutes the distinct haplotype, reflecting vicariant speciation since 1.55 Ma. However, this species was recently recorded from Yagachi-jima (Fig. 2 inset), northern Okinawa, and the haplotype is identical to the haplotype of S (southern) Iriomote-jima, suggesting recent long-distance dispersal. The Miyako and the original southern Okinawa populations (Fig. 2 inset) were included in the same Miyako haplotype network, and the southern Okinawa population originated from the Miyako population, dispersed at 0.3 Ma (Fig. 6). The network of the recently recorded specimens in southern and central Okinawa such as Tsuken (Fig. 2 inset) suggests recent dispersal also from the Miyako islands. Red double line: barrier for vicariance. Orange heavy line: *M. hebes* is separated by *M. minuta*, but genetically closely related.

**Fig. 9.**
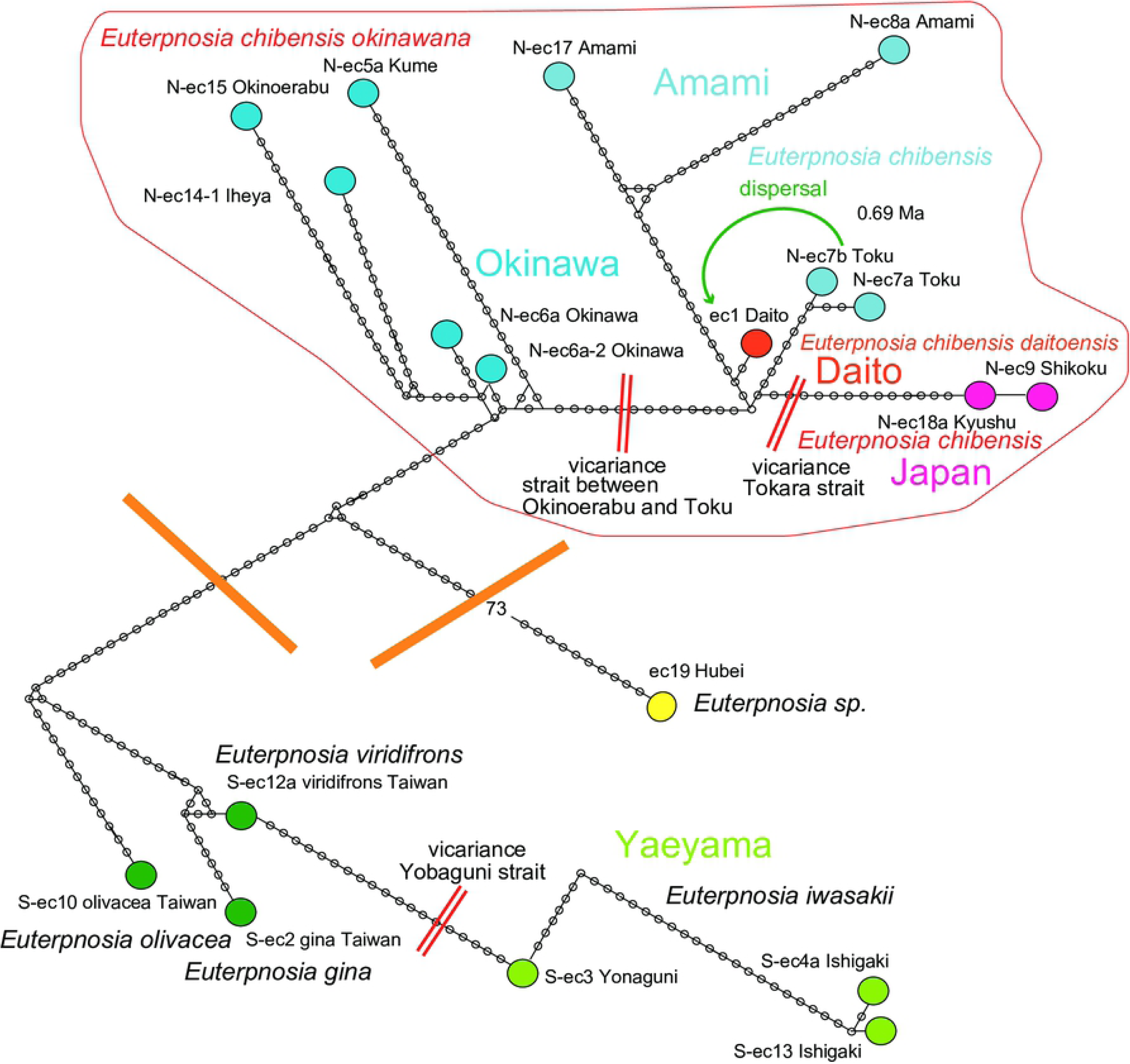
Haplotype network of *Euterpnosia.* The haplotypes were divided into *E. chibensis okinawana* (Okinawa) and *E. chibensis* networks, and the latter divided into the Amami and Japan networks, reflecting the 1.55 Ma event of vicariant speciation acted on the Ryukyu islands. However, *E. chibensis daitoensis* from the island of Daito was included in the Amami network, showing particularly close resemblance to the Toku network, suggesting that the Daito population was dispersed from the Tokuno-shima island at 0.69 Ma (Fig. 6).

**Fig. 10.**
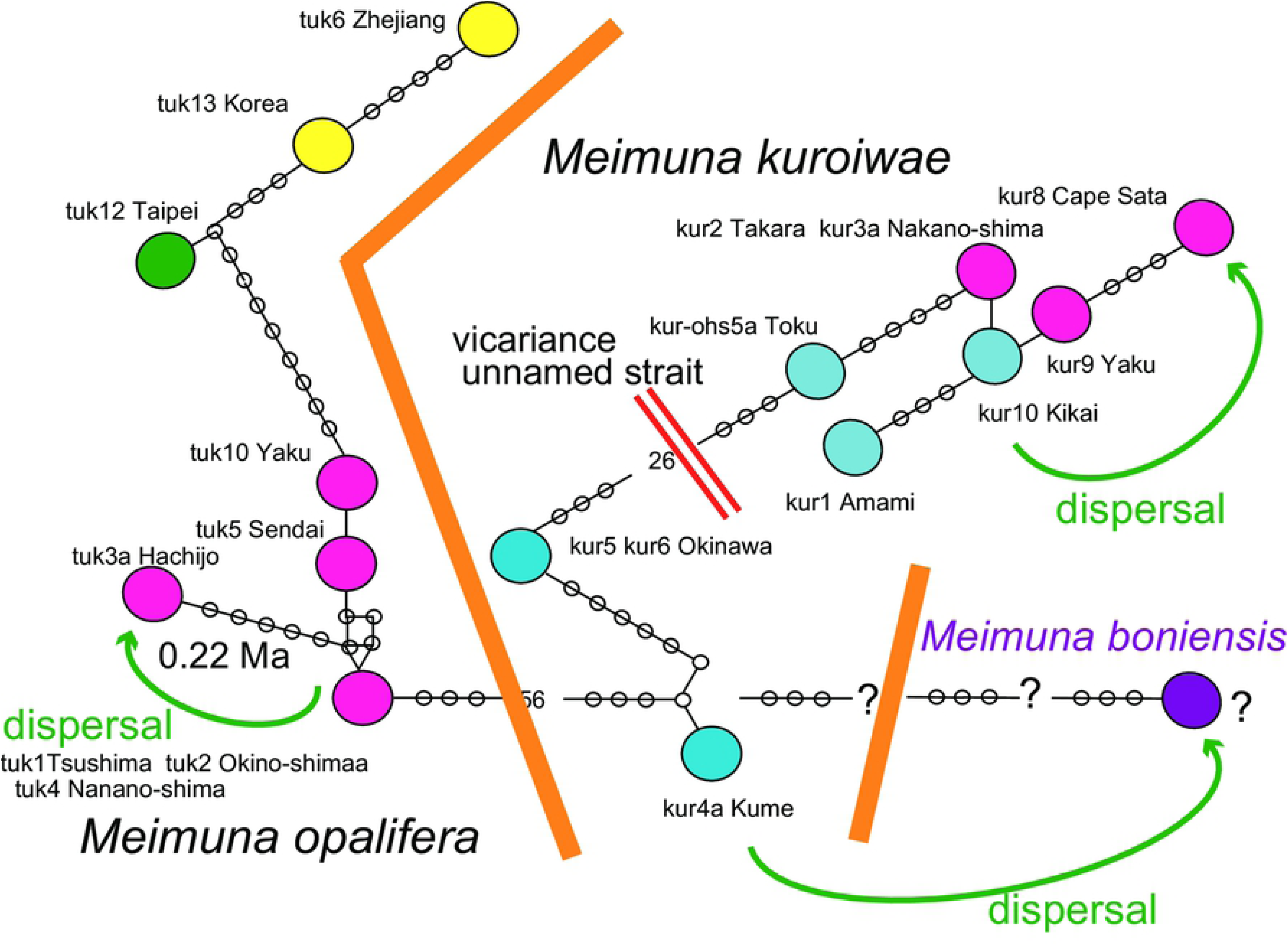
The haplotype network of *Meimuna. Meimuna kuroiwae* and *M. opalifera* constitute different networks. The *M. kuroiwae* network divided into the Okinawa and Amami-Tokara-Osumi-Kyushu networks, and the population of the southern end Kyushu may have been derived by dispersal from the southern islands. *Meimuna boniensis* may have been dispersed from Okinawa in ancient times (Nagata, 2019).

Because cicada nymphs live underground for two to five years (*Meimuna opalifera*: exceptionally one to two years; Hayashi & Saisho, 2011; c.f., *Magicicada*: 13-year and 17-year in eastern North America; e.g., Cooley et al., 2018), nymphs are unable to cross the seaways, so cicadas tend to be affected by vicariant speciation, as reflected in the vicariance of analyzed cicada species.

### Natural and artificial dispersals

#### Reason for dispersal

Some network patterns are explained by occasional dispersal, even where severe vicariance acted on these cicadas. Dispersal may be natural or artificial (anthropogenic), and modern or ancient for natural dispersal.

In the *Cryptotympana facialis* network (Fig. 7), the Japanese haplotype constitutes a part of the Yaeyama haplotype, and the mild differentiation between them is hard to explain by simple vicariance, in contrast to the case of the Yaeyama vs Miyako haplotypes. We suggest dispersal of the Japanese population from the Yaeyama population (green arrow in Fig. 7), and the dispersal date has an estimated node age of 0.7 Ma between the Japan and Yaeyama clades in the dated BI tree (Fig. 6). Note that the haplotype shows a simultaneous diffusional pattern (Fig. 7).

The original habitat of *Mogannia minuta* was on the Chinen peninsula of southern Okinawa (Fig. 2 inset), but it spread northward since 2004 (Sasaki, 2011; record in websites since 2012). The haplotype network consists of Yaeyama and Miyaki-Okinawa (Fig. 8), but sequence data or haplotype of min-mn9a Okinawa Yagachi (Fig. 2 inset) is the same as min-mn14a S Iriomote, and the Yagachi population might have dispersed from the southern Iriomote-jima before 2012 (Yagachi record is since 2012). Note that northern Irionote haplotype of min-mn4 N Iriomote differs by one base pair from S Iriomote (Fig. 8). The new habitat haplotype of min-mn6 Okinawa Tsuken (Fig. 2 inset) is similar to min-mn5 Miyako, and also dispersed from the Miyako islands before 2012, rather than from the original habitat on the Chinen peninsula. The Miyako islands consist of the Miyako-main, Shimoji, and Tarama islands. Whereas we did not analyze Shimoji and Tarama island specimens, we expect that they would be the same as min-mn6 Okinawa Tsuken. The haplotype of min-mn10a Okinawa Tamagusuku on the Chinen peninsula (Fig. 2 inset) is also similar to min-mn5 Miyako, although more differentiated than min-mn6 Okinawa Tsuken. *Mogannia minuta* originally of southern Okinawa was dispersed at about 0.3 Ma, estimated by the node age between the Miyako and the original Okinawa clades in the dated BI tree (Fig. 6).

The Daito oceanic islands were originally a remnant volcanic arc separated from the Izu volcanic arc by a series of back arc spreading events, including the formation Shikoku basin, concluding at 15 Ma. Thereafter the islands such as the Daito islands on the Philippine Sea plate have approached the Ryukyu continental arc, and have begun to collide with the trench/subduction zone or will collide in the future. On the Daito islands, far from the continental landmass, cicada may have originally been absent. Haplotype of ec1 Daito is close to N-ec7ab Toku (Fig. 9), and the Daito *Euterpnosia* population may have dispersed from the Tokuno-shima island. The dispersal date is an estimated node age of 0.69 Ma between the Daito and Toku clades in the dated BI tree (Fig. 6).

Hachijo oceanic island is a part of the Izu volcanic arc, its *Meimuna opalifera* haplotype is included in the Japanese network (Fig. 10), and should have dispersed from the Japan main islands. The dispersal date is an estimated node age of 0.22 Ma between the Hachijo and the resting Japan clades in the dated BI tree (Fig. 6). We recently estimated the emergent time of Hachijo as an island at 0.24 Ma (Osozawa et al., 2020). If the *Meimuna boniensis* haplotype is connected to the Okinawa haplotype, as Nagata (2019) inferred, the Bonin population may have dispersed from Okinawa, and the date was estimated at 1.4 Ma (Nagata, 2019).

### Dispersal by typhoon as a viable mechanism

#### Recent records and observations

Typhoon marginal winds rotate counter-clockwise at lower altitudes in the northern hemisphere, including Ryukyu islands, affected by Coriolis force. The marginal wind is strongest east of the eye, reinforced by the westerlies. In the eye, the warm rising air also rotates counter-clockwise, but radial flow from the eye at more than 10,000 m altitude is clockwise also affected by Coriolis force. A super typhoon is defined by an area of more than 800 km in radius affected by wind velocity of >15 m/s.

Flying insects can be carried by typhoon marginal winds. If the velocity is 15 m/s and distance between pick up and release points is 1000 km, the transport time is only 18.5 hours, and the insects may be able to survive during the travel.

A modern example of live insects transported by typhoon marginal wind was shown by Shoji (1991). *Polygonia c-aureum* (a nymphalid butterfly) is absent from the Ryukyu islands. However many specimens of its autumn form were collected as stray butterflies on Ishigaki, Iriomote, Hateruma, and Yonaguni islands starting in September 19, 1990. The butterflies were probably transported by southward marginal wind of the super typhoon 19th, 1990, from temperate Korea or northern China to these subtropical islands (Fig. 11). Typhoon generally tracks around the western margin of the Pacific High, northwestward in lower latitude affected by the trade wind, and then northeastward affected by the westerlies. The typhoon tends to be stationary at the turning point shown in Fig. 11.

**Fig. 11.**
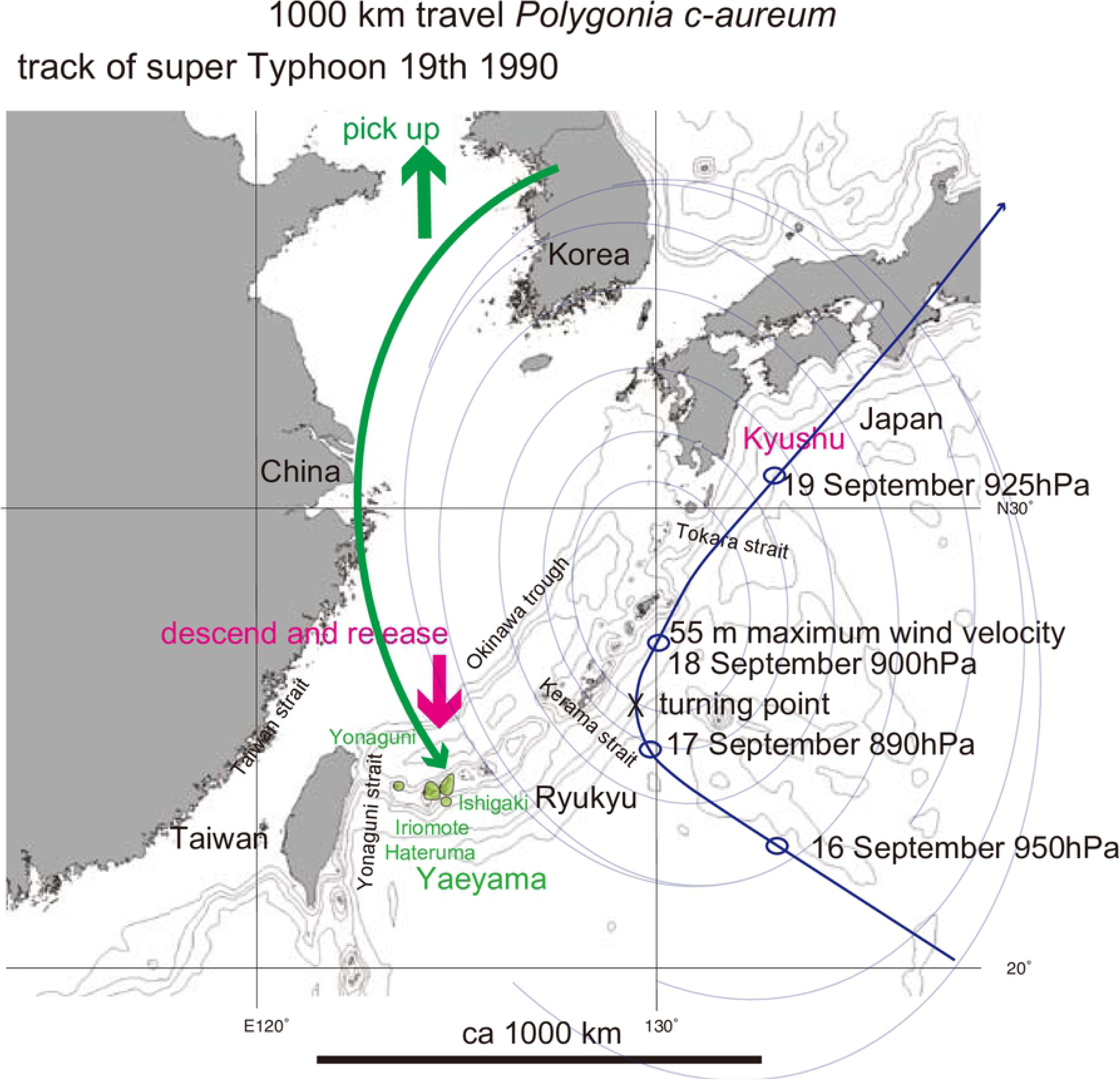
Track of super Typhoon 19th, 1990 (blue fine curved path), and 1000 km inferred transport of *Polygonia c-aureum* by typhoon winds (green curved arrow). The bold short arrows schematically (not to scale) show where the typhoon winds picked up cicada (green) and where the same cicada were released (red).

Flying insects can be carried in the raising air at the typhoon eye after transport by the marginal wind, and as the typhoon abates, the insects descend to the ground at this release point, but these may be somewhat exceptional cases.

One such exception was reported by Fukuda & Kanai (2013). Typhoon 17th, 2013, was generated offshore of northern Taiwan as a tropical cyclone at 2100 h on 31 August, migrated to the west of Yonaguni-jima, path along the west shore of the Ryukyu islands, landed on the Kagoshima (southernmost Kyushu) coast at 0300h on 4 September, diminished in strength and lost its eye over the Kagoshima city area, and declined to a temperate cyclone (Fig. 12). Many *Anax parthenope* were observed on Yonaguni-jima island on 2nd September but vanished on 3 September, probably carried away by the marginal winds of typhoon 17th, whose track was west of the Okinawa and then the Amami islands at these dates. Many active *A. parthenope* was later observed in the Kagoshima city area. More than 90 reports of this dragonfly mating and egg-laying were made by Kagoshima citizens to the Kagoshima Insect Society from the early morning of 4 September. These observation suggest that *A. parthenope* was transported by the typhoon 17th, with a distance over 1000 km from the Yonaguni-jima island to the Kagoshima city in 24 hours.

**Fig. 12.**
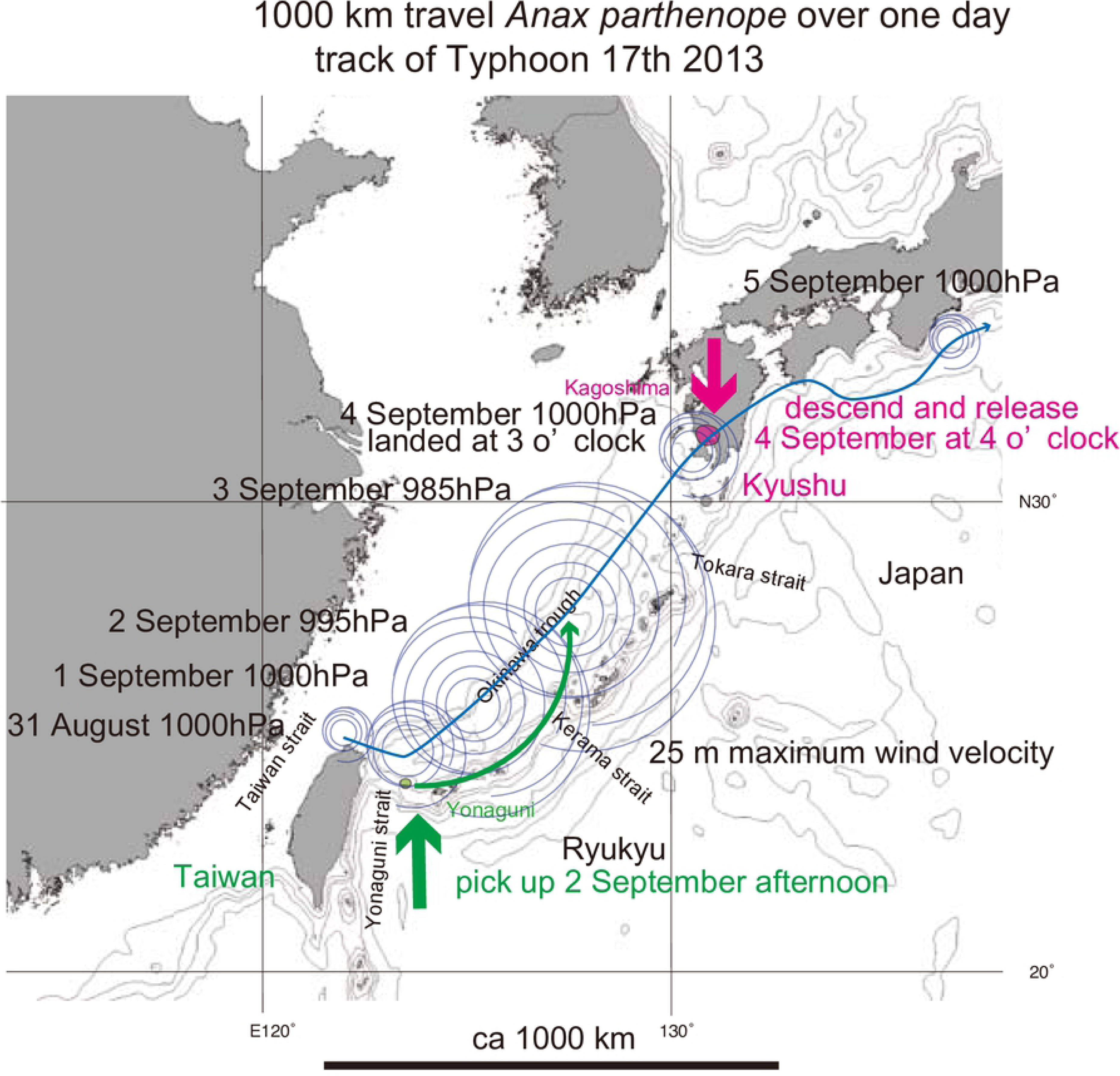
Track of Typhoon 17th, 2013 (fine blue sinuous path), and hypothetical 1000 km transport of *Anax parthenope* over one day by typhoon winds (green curved anticlockwise path). The bold short arrows schematically (not to scale) show where the typhoon winds picked up cicada (green) and where the same cicada were released (red).

Our phylogenetic study of another dragonfly, *Chlorogomphus* indicated recent migration from the Amami islands to Shikoku, Japan, as a result of transport by typhoon winds (Osozawa & Wakabayashi, 2015). Note that super typhoon are expected to have been generated in the East China Sea after 1.55 Ma, but not before that. This is because the sea floor spreading of that formed the Okinawa trough thereafter, typhoons tracks in this region would have made land fall and lost energy, rather than continuing northward to Japan (Osozawa & Wakabayashi, 2015).

#### Cryptotympana facialis

Counterclockwise marginal wind of super typhoon with a turning point in the mid East China Sea would have transported *Cryptotympana facialis* from Yaeyama to elsewhere Japan-Osumi-Tokara, passing through the Okinawa and Amami islands (Fig. 1; 0.7 Ma).

Owing to the large size of *C. facialis* compared to the minimal size of *Mogannia minuta*, one may consider whether this cicada can be safely transported for long distance before the release. The terminal settling velocity of tephra (volcanic ash) particles were estimated by considering Stokes’s law and Suzuki’s low with Cunningham correction (Shinbori, 2016). For a case of density = 1,000 kg/m^3^ (= water; cicada, ca. 700 kg/ m^3^), and radius of 10 mm (=*Cryptotympana facialis*; 1/3 for *Mogannia minuta*), the settling velocity is less than 10 m/s (7 m/s for the former; 3.3 x 0.7 = 2.3 m/s for the latter), but may be much slower considering the irregular shape and large wings of cicadas compared to idealized spherical or cubic shapes. Compared to the lateral air transport for super typhoon more than 15 m/s, cicadas, like violcanic ashes, may be able to drift in air and fly long distance before landing.

#### Mogannia minuta

Considering the sighting of this species at its dispersed location in 2004 (Sasaki, 2011) and the 2-year nymph period, the typhoon that transported mated female(s) was probably pre-2002, and considering the emergent season, the track time should have been May to July. Annual typhoon tracks are recorded on the Japan Meteorological Agency (https://www.jma.go.jp/jma/indexe.html) and related websites. Using these records we may evaluate the possible typhoons that transported the cicada(s).

Only Typhoon 5th, of early June 2002 (Fig. 13) appears viable to have transported the cicada by marginal wind. For comparison, of potential post-2004 tracks, only Typhoon 5th, late June 2011 (Fig. 13) had a track route with the potential for a similar marginal wind dispersal for *M. minuta*. That specific typhoon had the potential to disperse from Iriomote-jima as the Yagachi population found after 2012, but not the other Okinawa population dispersed from the Miyako islands (Fig. 2 inset), because late June is still the adult season on Iriomote-jima but not on the Miyako islands.

**Fig. 13.**
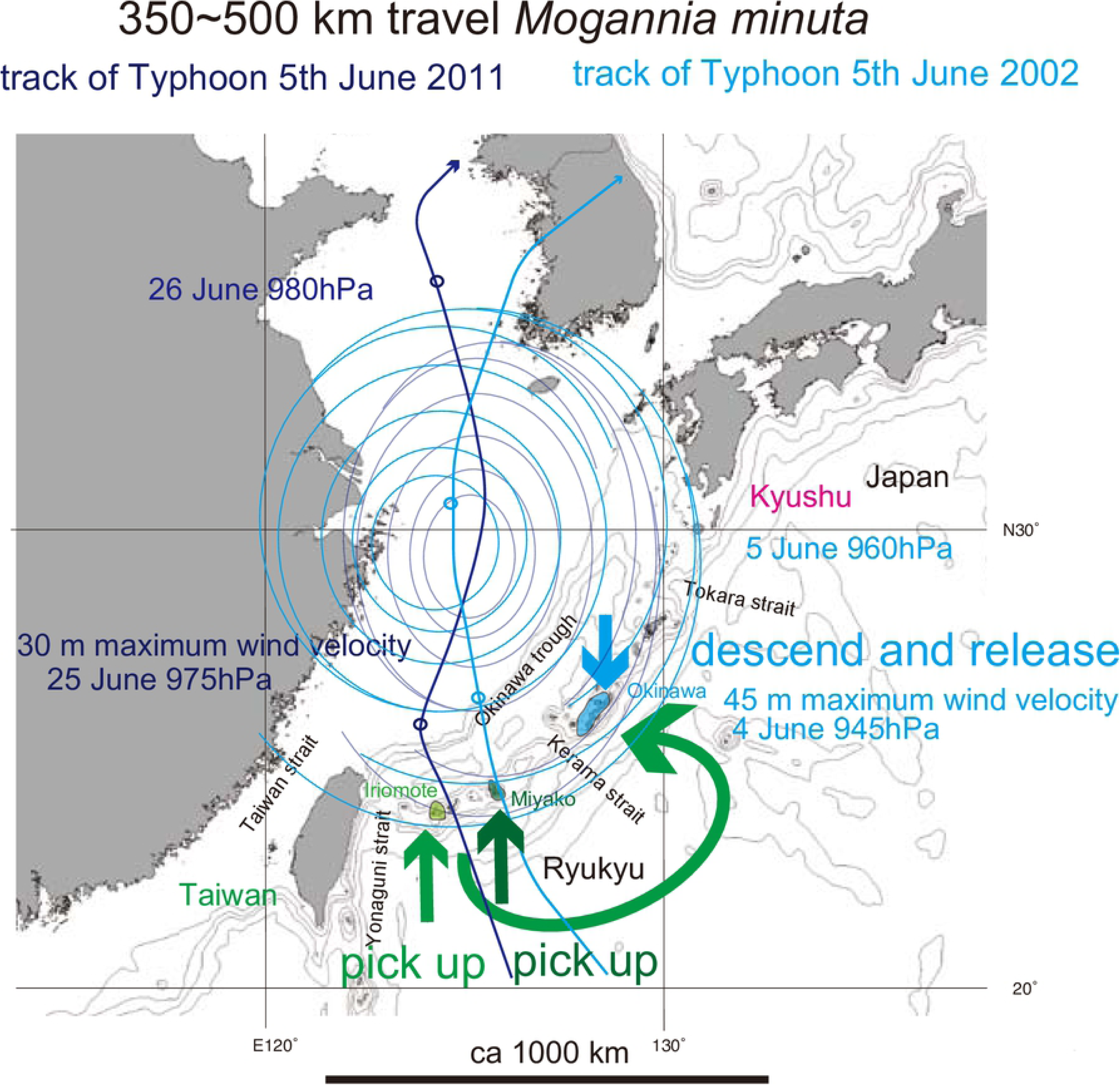
Track of track of Typhoon 5th, June 2002 (medium blue curved path) and track of Typhoon 5th, June 2011 (very dark blue sinuous path), and hypothetical 350∼500 km transport *Mogannia minuta* by the marginal typhoon winds (green curved path). The bold short arrows schematically (not to scale) show where the typhoon winds picked up cicada (green and dark green) and where the same cicada were released (blue).

Typhoon 1st, May 2001 (Fig. 14), that passed along the southern Ryukyu islands, over the Yaeyama and then Miyako islands on 13 May, lost energy east of Okinawa island on 14 May, and may have released the cicada. In Fig. 14, we also show the track of Typhoon 8th May 1997, another potential typhoon that may have transported *M. minuta* in its marginal winds.

**Fig. 14.**
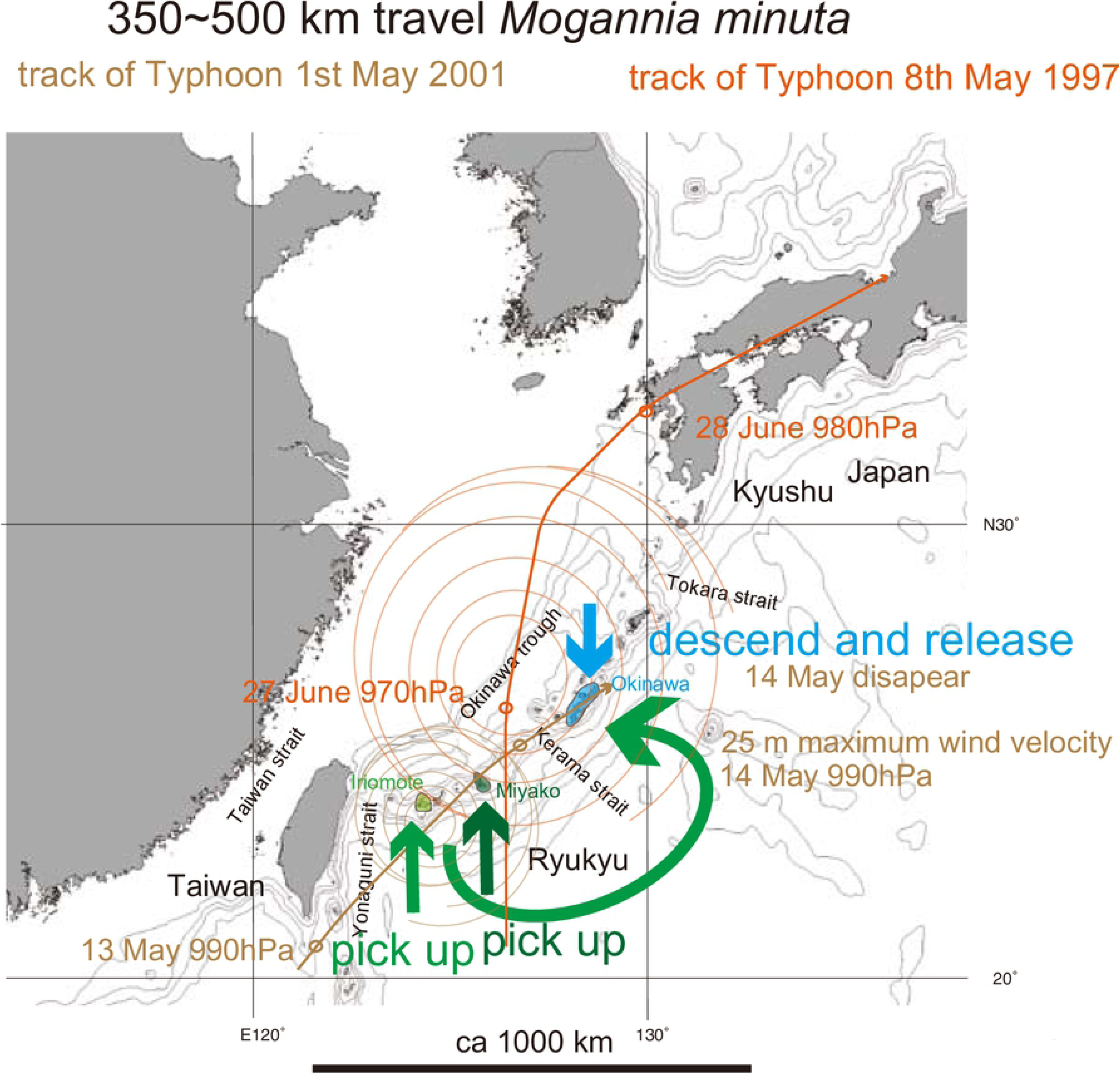
Track of Typhoon 1st, May 2001 (brown curved path), that may have transported and then released *Mogannia minuta* on 14 May when it was lost energy west of Okinawa. The track of Typhoon 8th, May 1997 (orange curved path), is also shown, because this typhoon also followed a path that may have transported *M. minuta* from the southern Ryukyu islands to Okinawa. The green path depicts this hypothetical transport. The bold short arrows schematically (not to scale) show where the typhoon winds picked up cicada (green and dark green) and where the same cicada were released (blue).

*Mogannia minuta* in the original habitat of southern Okinawa (Fig. 2 inset) may have been dispersed 350 km from Miyako at 0.3 Ma (Fig. 6) by marginal wind of typhoon with a turning point in the southern East China Sea (Fig. 2), or by a track similar to that of Typhoon 1st, May 2001 (Fig. 14).

#### Euterpnosia chibensis daitoensis

This endemic species on the Daito islands is a sister of *Euterpnosia chibensis* on Tokuno-shima island, and may have dispersed from Tokuno-shima at 0.71 Ma (Fig. 6). If a turning point of super typhoon was in the central Philippine Sea, the southeastward marginal wind west of the center could transport *Euterpnosia* of Toku toward Daito (Fig. 3).

*Euterpnosia chibensis* on the O-shima island of the northernmost Izu islands might have recently (pre 1998) dispersed from Honshu. Typhoon 5th, July 1996 may have transported *E. chibensis* on the Boso peninsula (Honshu) toward O-shima (Fig. 3).

#### Meimuna

*Meimuna boniensis* endemic on the Bonin islands is a sister of *M. kuroiwae* on the continental Okinawa islands (Nagata, 2019), and should have dispersed from Okinawa in pre-historic time, crossing the Pacific ocean. If a turning point of super typhoon was offshore southwest Japan, the eastward marginal wind south of the center may have transported *Meimuna* of Okinawa toward the Bonin islands (Fig. 4).

*Meimuna opalifera* habitat extends southward to the Hachijo and Aoga-shima islands, of the central Izu islands (Fig. 5). Note that habitat of *Cryptotympana facialis* is limited to Miyake island (Fig. 1), suggesting less flying ability for *C. facialis*. The southward dispersal at 0.22 Ma (Fig. 6) could be by southward marginal wind of typhoon offshore the Boso peninsula (Fig. 5).

*Meimuna opalifera* is also known from the northern Tokara islands, but also on the uninhabited Yokoate island, south of the Tokara strait (Fig. 5). It is absent on the southern Tokara islands of Takara and Kodakara, so the Yokoate population may have been dispersed from the northern Tokara islands by transport by typhoon marginal winds.

### Rafting of egg-bearing plants as another possible dispersal mechanism

The egg stage is only 40 days for *Mogannia minuta* (Hayashi, 1976), whereas it is close to one year for other cicadas (Hayashi & Saiso, 2011). Female cicadas generally lay eggs in dead, decaying wood, but *M. minuta* deposits eggs on sugarcane leaves (Hayashi & Saiso, 2011). Such material (wood or leaves) may be rafted on the Kuroshio current (Fig. 1), and eggs can be transported from a southerly island to a more northerly one.

We suggested that *Carabus blaptoides* beetles from central Kyushu were carried on driftwood by a major flood on 11–14 July 2012 to the ocean, then transported by rafting on the Kuroshio current to eventual landfall on Nii-jima islet, of the northern Izu islands (Osozawa K et al., 2016). The rafting was taken only ten days, and the beetle can apparently survive that duration of rafting.

For *Cryptotympana facialis*, rafting of eggs on/in driftwood on the Kuroshio current from Yaeyama especially to Tokara (Fig. 1) at 0.7 Ma (Fig. 6) may have been possible. Similar transport of *Meimuna boniensis* by rafting on the Kuroshio and then Ogasawara currents from Okinawa, through the Izu islands, to the Bonin islands (Figs. 4 and 10) may have possible. *Meimuna kuroiwae* may have potentially rafted on the Kuroshio current from the Amami islands to Cape Sata (Figs. 4 and 10) at 0.43 Ma (Fig. 6).

There are several problems with rafting as a cicada dispersal mechanism. The first concerns the derivation of the driftwood from the original island, although the above are lowland and coastal species (Fig. 1). Survival of eggs in seawater for at least ten days is also uncertain. A final problem is the vulnerability of hatched nymphs on material washed up on a beach. They may have to crawl a fairly long distance to reach the forested cover where they would burrow. Owing to these issues, we suspect that typhoon dispersal may be more likely than rafting dispersal for cicadas.

### Artificial dispersal

For *Cryptotympana facialis*, human transplanting, of nymphs within the soils around roots or branches with eggs was proposed for the mechanism of their invasion into the Amami islands from the Okinawa islands (Fukuda & Morikawa, 1999). We revisit their proposal, because paper was published in a journal of limited circulation and in Japanese.

*Cryptotympana facialis* was not recorded in the Amami islands (Fig. 1) from at least 1959 until 1990 (Fukuda, 1991ab). However, the first record of 1991 was from the Seisui park, Setouchi, southern Amami Oshima (Tahata, 1994 cited in Fukuda et al., 2006). None of the recorded sightings in 1991 by Fukuda (1991b) was from northern Amami Oshima.

A modern ferry from Okinawa, Toku to Amami began service in 1972, and a ferry from Amami to Kikai-jima started service in 1974, whereas there has been no direct ferry from Okinawa to Kikai-jima (Fig. 1). The development of these routes may have been related to the return of Okinawa from the USA in 1972, and the following economic development funded by the Japan government. On the Amami islands, ports, airports, roads, public facilities including urban parks were built, and extensive landscaping along shores and valleys was undertaken. Parks were planted with trees, and flowering trees such as *Erythrina variegata* were imported from Okinawa by the ferries connected with Amami. *C. facialis* on the Amami islands was first recorded at or near newly constructed urban parks in low-lying areas, several years after Okinawa trees were planted. Because cicada nymphs are usually underground for several years, and eggs hatch the next year, so a lag time between the actual transplanting and adult emergence may have been several years. Table 2 summarized dates of the first collection of *C. facialis*, and the coincidence of the planting and the first adult emergence dates supports the hypothesis of the artificial dispersals from Okinawa at least for Amami Oshima.

**Table 2.** Summary of dates of first records of *Cryptotympana facialis* and transplantation dates for the Amami islands after references shown on this table. For land formation (human landscaping) dates, documentation by archival aerial photos offered by Geospatial Information Authority of Japan are referenced.

As pointed out by Fukuda & Morikawa (1999) and Fukuda et al. (2006), the Amami population lacks the abdominal white band, as is the case for the Okinawa population. The present study showed that COI sequence of most of the Amami and Toku populations except for the Setouchi, a part of southern Amami population, is common to the sequence Okinawa population. These are concordant to the above observation summarized in Table 2 and their explanation.

For *Mogannia minuta*, that lays eggs on sugar cane, transplanting with nymphs and eggs from Miyako-jima or Iriomote-jima may be expected to result from modern shipping. However, sugar cane harvesting is mostly by cutting (rather than uprooting), and the import from another island is not necessary owing to self-sufficient crops. In addition, sugar cane and the root is lost after the harvest from the field, and nymphs usually cannot survive after the harvest. In contrast, the habitat for *Miscanthus sinensis* is adjacent to the sugar cane field, and we observed and collected *M. minuta* there.

### Alternative proposal for absence of *Cryptotympana facialis* on the Amami islands

In addition to long distance dispersal by typhoon winds, we need to discuss the genetic homogeneity with a uniform COI sequence covering an extensive region that includes the Japanese, Osumi, and Tokara islands and encompassing more than 100 specimens, except for 4 specimens differing by one base substitution (Fig. 1). After the dispersal to locations within the Japanese, Osumi, and Tokara islands at 0.7 Ma (Fig. 6), *C. facialis* should have extensively radiated and expanded from original released positions. The genetic similarity may reflect bottleneck or founder effect, and the following range expansion.

The above consideration suggested that wide range of *Cryptotympana facialis* in the Japan, Osumi, and Tokara islands developed after 0.7 Ma. Accordingly, the Amami islands south of the Tokara islands may have originally lacked this species before 0.7 Ma.

We propose, however, an alternate explanation for why *Cryptotympana facialis* was absent in the Amami islands prior to 1991. This species inhabits coastal lowlands less than 100 m in altitude in the Ryukyu islands including the post-1991 habitat on the Amami islands. The Toku-Kikai islands and Okinoerabu island have a distinct uplift history (Fig. 1; Osozawa et al., 2012 and modified). The Toku-Kikai islands have three marine terraces recording uplift events at 0.9, 0.4 or 0.2 Ma, and relatively recent (Figs. 15 and 16) or alternatively reflecting relatively fast progressive uplift with erosion of wavecut benches (future terraces) during high stands of sea level. The uplift of these islands is an apparent consequence of the collision of the Amami and Daito plateaus with the west-dipping subduction zone that passes beneath the Toku-Kikai islands. On the Toku-Kikai islands this uplift history has resulted in a landscape lacking in lowland habitat that features seacliffs or terrace-riser cliffs (cliffs bounding marine terraces) instead (Figs. 15 and 16). As a result, had there been an original *C. facialis* population, it may have been eliminated by tectonic (geologic) destruction of habitat on the Toku-Kikai islands. In contrast, Amami Oshima, except for its northeastern end with a terrace, has subsided so that upland areas rise from the coast instead of having the more gentle topography that typifies the ideal lowland habitat of *C. facialis*. Although Okinoerabu-jima has been uplifted, the landscape evolution is different than the Toku-Kikai islands that have concentric marine terraces and associated cliffs. Okinoerabu-jima has a wide lowland region consisting by a single terrace, and lacks sea cliffs except for the western part of the northern shore where there is a sea cliff that has resulted from an active fault scarp. The Okinoerabu geomorphic features have formed since the inception of uplift at 1.55 Ma and likely reflect a lower rate of uplift compared to Toku-shima and Kikai-jima (Fig. 17).

**Fig. 15.**
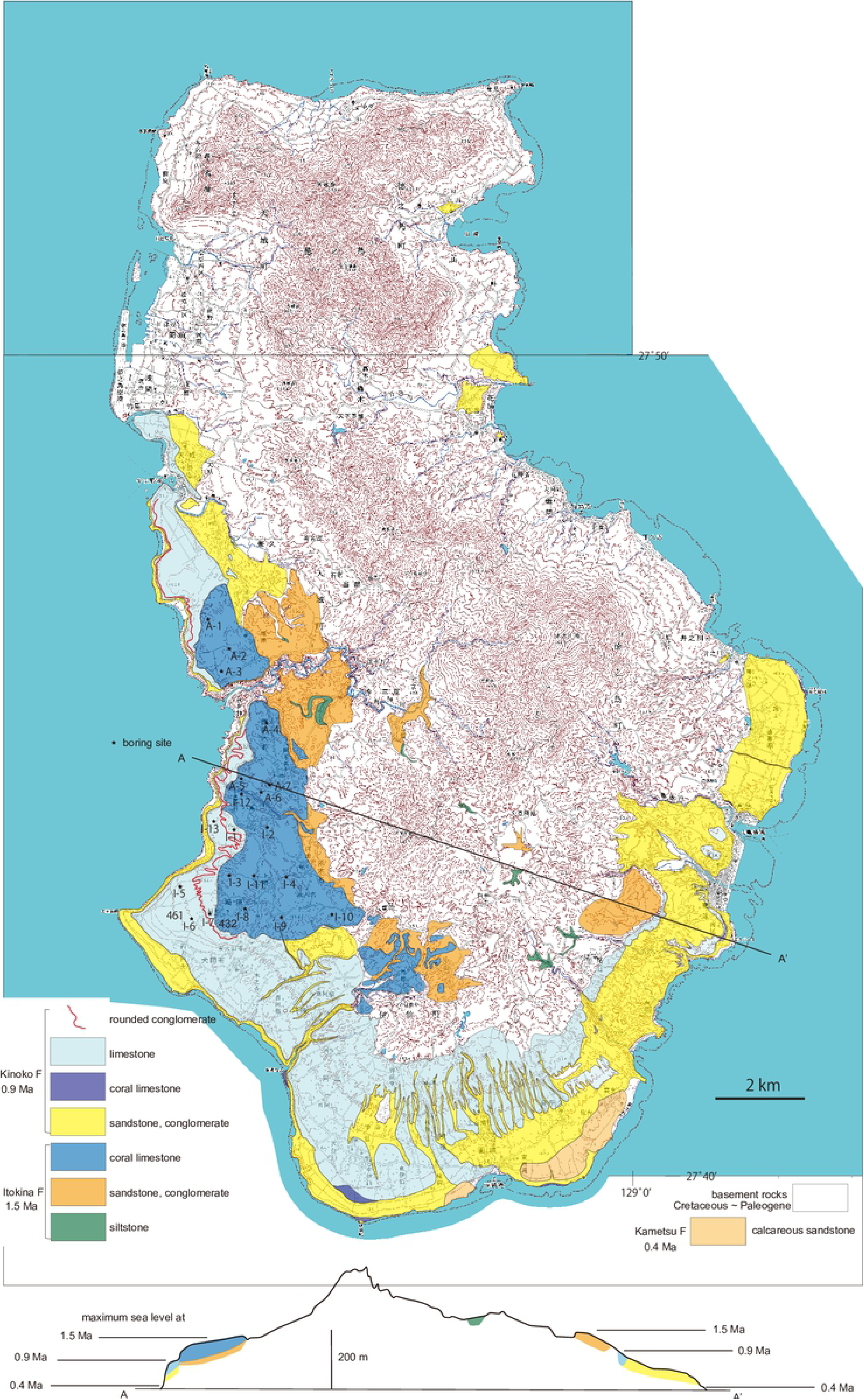
Geological map and cross section (below) of Tokuno-shima island. Three terraces consisting of the 1.5, 0.9 and 0.4 Ma limestone show three episodes of uplift or progressive uplift with erosion of wavecut benches during high sea level stands. Note the lack of coastal plains at 0.9 and 0.4 Ma, with development of 100-m-high sea cliffs instead.

**Fig. 16.**
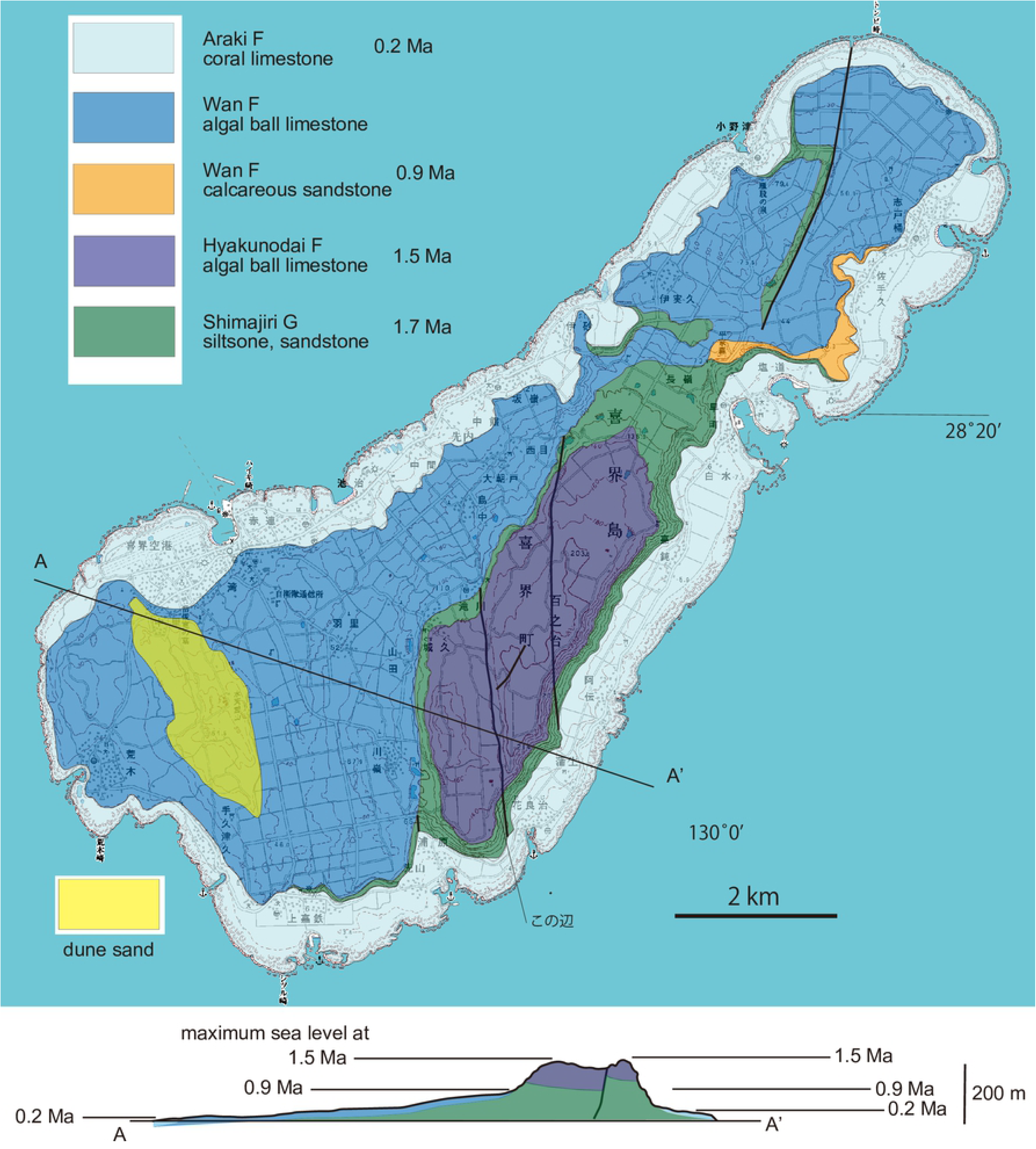
Geological map and cross section (below) of Kikai-jima island. Three terraces of the 1.5, 0.9 and 0.2 Ma limestone show three episodes of uplift or progressive uplift with erosion of wavecut benches during high sea level stands. Note the lack of a coastal plain at 0.9 Ma (100 m high sea cliff), but a relatively wide coastal plain at 0.2 Ma above a lower sea cliff.

**Fig. 17.**
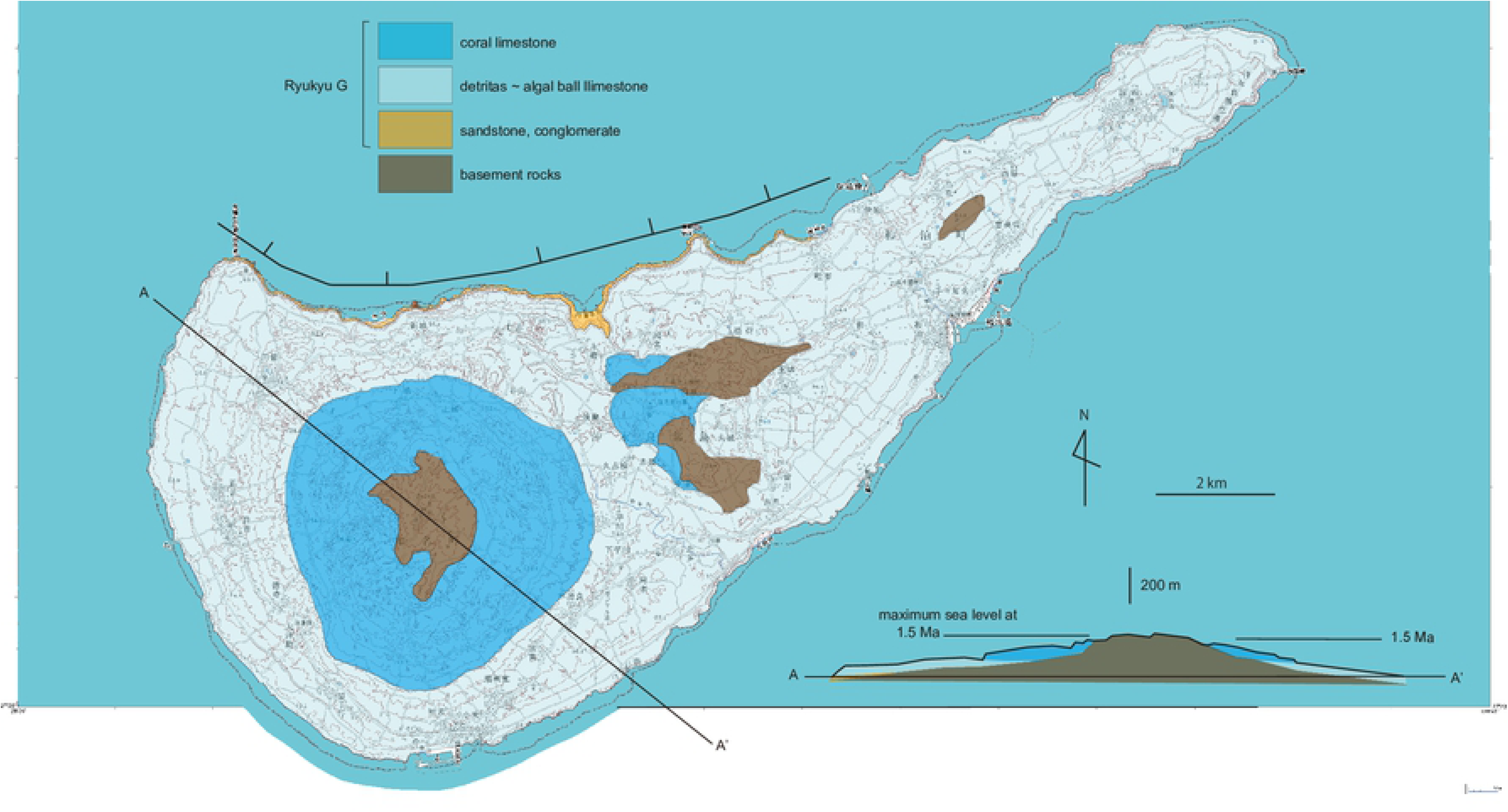
Geological map and cross section (below) of Okunoerabu-jima. A terrace developed on 1.5 Ma limestone. Uplifting was apparently slower compared to the examples of Toku-shima and Kikai-jima and has been progressed since 1.55 Ma. This history has resulted in a relatively wide coastal plain lacking high sea cliffs, except for fault scarp cliff at along the western part of northern margin. Miyako-jima and Ishigaki-jima have also only a single terrace in 1.5 Ma limestone, and northeastern end of Amami Oshima is also associated with 1.5 Ma limestone, whereas Okinawa-jima and Kume-jima have low elevatioon terraces developed with ca. 0.2 ∼ 0.4 Ma (not dated) limestone.

Accordingly, the lack of *C facialis* in the Amami islands prior to 1991 introduction may have been a consequence of the species not previously inhabiting the islands, or the species may have once populated parts of these islands but vanished when its habitat was destroyed by vertical crustal movements associated with the tectonics of this Amami region, especially Toku-Kikai.

## Conclusions

*Cryptotympana*, *Euterpnosia*, *Mogannia*, and *Meimuna* cicadas in the Ryukyu islands and the other parts of east Asia experienced the vicariant speciation due to the physical isolation of these islands from the Chinese continent since 1.55 Ma (Quaternary), and this tectonic and related evolutionary process continues in this region. Some cicadas were, however, transported distances of up to more than 1000 km in both recent (historic) and ancient (pre-historic) times. Winds associated with super typhoons that approached and passed through the Ryukyu islands may have been the key agent of the long-distance cicada dispersal. Because area of the present East China Sea was east Asian Chinese mainland before 1.55 Ma, typhoons would have lost energy upon landfall. In contrast, after the formation of the East China Sea by back-arc rifting (sea-floor spreading) starting at 1.55 Ma, typhoons would amplify or maintain their strength in this region.

## Declaration of competing interest

The authors declare that they have no conflict of interest.

## Supplementary Material

Supplementary data in Table 1 are available at GenBank / DDBJ.

## Data Availability Statement

All relevant data are within the manuscript.

## Acknowledgements

Tutorial of Network 10 was by Hiroaki Kaneko. We thank cicada collectors shown in Table 1 for offering specimens available for our analyses. Bor-ming Jahn (Taiwan University; deceased 1 December, 2016), Ping-Shih Yang (Taiwan University), Chin-Ho Tsai (National Dong Hwa University), and Jen-Zon Ho and Hua-Te Fang (Endemic Species Research Institute) supported sample collections and obtained permission to collect in Taiwan.

## Funding

This project was partly financed through the Osozawa Fund (Former), Tohoku University. We thank Keiji Nunohara (Nunohara Office for Geological Survey), Kohei Sugawara (Ecofarm GSK), CTI Engineering Co., Ltd., and NEWJEC, Inc. for contributing to this fund.

## Author Contributions

Soichi Osozawa collected samples and coordinated the research, carried out the DNA analyses, and wrote this paper, Kenichi Kanai and Haruo Hukuda collected samples and supervised, and John Wakabayashi (native English-speaking American geologist) edited the writing.

